# Neuronal post-developmentally acting SAX-7S/L1CAM can function as cleaved fragments to maintain neuronal architecture in *C. elegans*

**DOI:** 10.1101/2021.01.25.428117

**Authors:** V.E. Desse, C.R. Blanchette, P. Perrat, C.Y. Bénard

**Affiliations:** Université du Québec à Montréal, Department of Biological Sciences, 141 President Kennedy Avenue, Montréal, QC H2X 1Y4, Canada; University of Massachusetts Medical School, Department of Neurobiology, 364 Plantation St, Worcester, MA 01605, USA

**Keywords:** neuronal maintenance, lifelong, L1, *sax-7*, Ig, cleavage

## Abstract

Whereas remarkable advances have uncovered mechanisms that drive nervous system assembly, the processes responsible for the lifelong maintenance of nervous system architecture remain poorly understood. Subsequent to its establishment during embryogenesis, neuronal architecture is maintained throughout life in the face of the animal’s growth, maturation processes, the addition of new neurons, body movements, and aging. The *C. elegans* protein SAX-7, homologous to the vertebrate L1 protein family, is required for maintaining the organization of neuronal ganglia and fascicles after their successful initial embryonic development. To dissect the function of *sax-7* in neuronal maintenance, we generated a null allele and *sax-7S*-isoform-specific alleles. We find that the null *sax-7(qv30)* is, in some contexts, more severe than previously described mutant alleles, and that the loss of *sax-7S* largely phenocopies the null, consistent with *sax-7S* being the key isoform in neuronal maintenance. Using a sfGFP::SAX-7S knock-in, we observe *sax-7S* to be predominantly expressed across the nervous system, from embryogenesis to adulthood. Yet, its role in maintaining neuronal organization is ensured by post-developmentally acting SAX-7S, as larval transgenic *sax-7S*(+) expression alone is sufficient to profoundly rescue the null mutants’ neuronal maintenance defects. Moreover, the majority of the protein SAX-7 appears to be cleaved, and we show that these cleaved SAX-7S fragments together, not individually, can fully support neuronal maintenance. These findings contribute to our understanding of the role of the conserved protein SAX-7/L1CAM in long-term neuronal maintenance, and may help decipher processes that go awry in some neurodegenerative conditions.

## Introduction

An important yet poorly understood question of neurobiology is how the organization of neural circuits is maintained over a lifetime to ensure their proper function. Largely established during embryogenesis, the architecture of the nervous system needs to persist throughout life in the face of the animal’s growth, the addition of new neurons, maturation processes, body movements, and aging. Whereas significant progress has been made in understanding the processes driving neuronal development, little is known about the mechanisms ensuring lifelong maintenance of nervous system architecture and function.

Research using *C. elegans* has uncovered a number of immunoglobulin (Ig) superfamily molecules required for the long-term maintenance of neuronal architecture (Benard and Hobert, 2009). These include the large extracellular protein DIG-1 (Benard et al., 2006; Johnson and Kramer, 2012), the small two-Ig domain proteins ZIG-3, ZIG-4, and ZIG-10 (Aurelio et al., 2002; Benard and Hobert, 2009; Benard et al., 2012; Cherra and Jin, 2016), the ectodomain of the FGF receptor EGL-15 (Bülow et al., 2004), as well as SAX-7 (Pocock et al., 2008; Sasakura et al., 2005; Wang et al., 2005; Zallen et al., 1999; Zhou et al., 2008). Here, we further the investigation of SAX-7/L1CAM’s role in the lifelong maintenance of neuronal architecture.

SAX-7 is an evolutionary conserved transmembrane cell adhesion molecule homologous to mammalian L1CAM (Chen et al., 2001; Hortsch, 2000; Hortsch et al., 2014). In *C. elegans*, SAX-7 exists as two main isoforms, a long isoform SAX-7L and a short isoform SAX-7S. These two isoforms are identical for their intracellular tail, transmembrane domain (TM), and most of their extracellular region including five identical fibronectin type III domains (FnIII), and four Ig-like domains. They differ in the N-terminal extracellular region, where SAX-7S has four Ig domains (Ig 3-6), whereas SAX-7L has six Ig domains (Ig 1-6). Transgenes of SAX-7S, but not of SAX-7L, rescue the defects of *sax-7* loss-of-function mutants, indicating that the SAX-7S isoform is central to *sax-7* functions (Pocock et al., 2008; Ramirez-Suarez et al., 2019; Sasakura et al., 2005; Wang et al., 2005). Vertebrate proteins of the SAX-7/L1CAM family include L1CAM, NrCAM, CHL1, and Neurofascin (Brummendorf et al., 1998; Brummendorf and Rathjen, 1996; Haspel and Grumet, 2003; Hortsch, 2000; Hortsch et al., 2014).

*sax-7*/L1CAM is well known to contribute to the development of distinct neurons in *C. elegans*. It is involved in dendrite development and axon guidance (Cebul et al., 2020; Chen et al., 2019; Diaz-Balzac et al., 2015; Diaz-Balzac et al., 2016; Dong et al., 2013; Heiman and Pallanck, 2011; Ramirez-Suarez et al., 2019; Salzberg et al., 2013; Schafer and Frotscher, 2012; Sherry et al., 2020; Yip and Heiman, 2018; Zhao et al., 1998; Zhu et al., 2017). In flies and mammals, homologues of *sax-7* function in neuronal migration, axon guidance, and synaptogenesis (Bieber et al., 1989; Godenschwege et al., 2006; Hall and Bieber, 1997; Rougon and Hobert, 2003; Sonderegger et al., 1998). In humans, mutations in L1CAM severely impair neuronal development, leading to disorders collectively referred to L1 or CRASH syndrome for corpus callosum hypoplasia, mental retardation, aphasia, spastic paraplegia and hydrocephalus (Fransen et al., 1997; Hortsch et al., 2014).

Besides their roles in neuronal development, SAX-7/L1CAM family members also function in the mature nervous system to preserve neuronal organization. In *C. elegans, sax-7* is required for maintaining neuronal organization well after development is completed, as specific neuronal structures that initially develop normally in *sax-7* mutant animals, later become disorganized. For instance, in *sax-7* mutants, a subset of axons within the ventral nerve cord, which developed normally during embryogenesis, become displaced to the contralateral fascicle during the first larval stage; and neurons within embryonically established ganglia become progressively disorganized by late larval stages and adulthood in *sax-7* mutants (Pocock et al., 2008; Sasakura et al., 2005; Wang et al., 2005; Zallen et al., 1999; Zhou et al., 2008). Such post-developmental neuronal disorganization displayed by *sax-7* mutant animals can be prevented if animals are paralyzed (Pocock et al., 2008; Sasakura et al., 2005), indicating that the mechanical stress from body movements contributes to perturbing neuronal architecture in these mutants. In mammals, roles for L1 family members in the adult nervous system have been revealed as well through the study of conditional knockouts. Adult-specific knockout of neurofascin affects rats behavior and alters the axon initial segment in mice (Kriebel et al., 2011; Zonta et al., 2011); knockout of L1CAM specifically in the adult mouse brain leads to behavioral deficits and synaptic transmission changes (Law et al., 2003); and CHL1 conditional depletion in a subtype of forebrain neurons in mice leads to defects in working memory duration (Kolata et al., 2008). Thus, L1CAM family proteins contribute to preserving the functionality of the mammalian adult nervous system.

Despite the evolutionarily conserved importance of SAX-7/L1CAM, its role in the long-term maintenance of the neuronal architecture remains unclear. In order to better understand how SAX-7/L1CAM participates in neuronal maintenance, here we have generated and characterized a null allele of *sax-7*, tested the temporal requirements for *sax-7S* neuronal maintenance function, determined the endogenous expression pattern of SAX-7S, and assessed the function of SAX-7S cleavage products in neuronal maintenance. Our results further our understanding of the roles of the evolutionarily conserved molecule SAX-7/L1CAM in the lifelong persistence of neuronal organization and function.

## RESULTS

### Molecular analysis of previous *sax-7* mutant alleles

The interpretation of previous structure-function analyses for *sax-7* was limited by the lack of a clear null mutation for the gene. In particular, the existence of gene product in *sax-*7(*nj48*), an allele reported to be a complete loss-of-function of the gene *sax-7*, has not been fully assessed. We examined *sax-7* transcripts by RT-PCR for *nj48*, as well as for other *sax-7* mutant alleles, including the *sax-7L*-specific alleles *eq2* and *nj53*, and two alleles that affect both *sax-7* isoforms, *tm1448* and *eq1* (**Fig. 1A,B**). We detected transcripts corresponding to all isoforms of *sax-7* in all mutants tested (**Fig. 1C**; all RT-PCR products were verified by sequencing), except when the primer targets a sequence that is deleted by a given mutation. In particular, transcript was detected in *nj48* mutants, using four different primer pairs (**Fig. 1C**), indicating that *nj48* is not a null allele.

**Fig 1.**
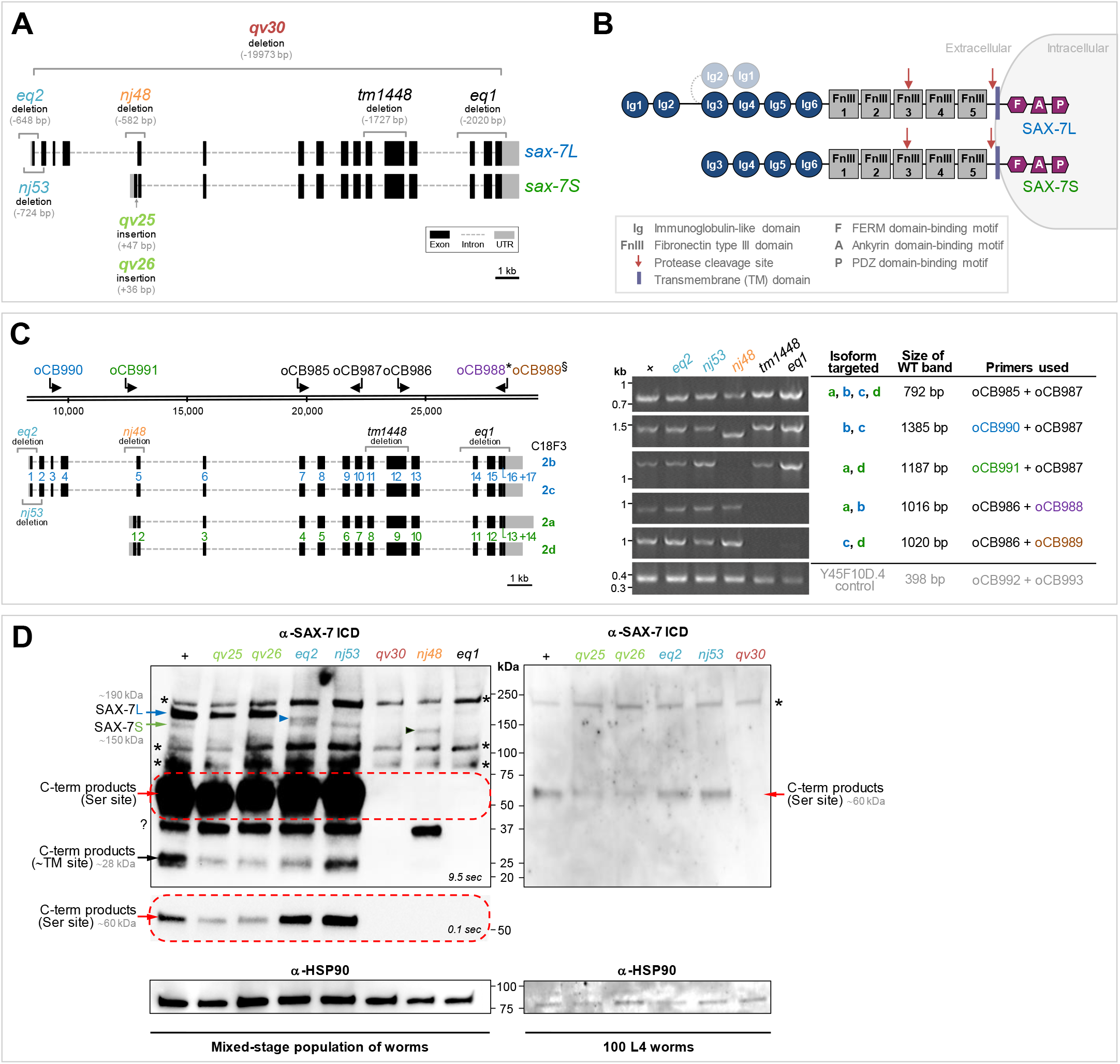
Analysis of *sax-7* mutant alleles. **(A)** Schematics of the gene structure for the *sax-7* short (C18F3.2a) and long (C18F3.2b) isoforms. The mutant alleles used in this study are indicated, including the newly generated the null allele *qv30* and *sax-7S*-specific alleles *qv25* and *qv26* (see **Fig. S1 A-C** for sequence information). Alleles *nj48, tm1448*, and *eq1* affect both isoforms, and alleles *eq2* and *nj53* are *sax-7L*-specific (see **Table 1** for allele information). **(B)** Schematics of the protein structure of SAX-7L and SAX-7S. Red arrows indicate cleavage sites: serine protease cleavage site in FnIII#3, or cleavage site proximal to the transmembrane (TM) domain. The two N-terminal Ig domains Ig1 and Ig2 may fold at the hinge region onto Ig3 and Ig4, indicated in grey (Pocock et al., 2008). **(C)** Schematics of the four encoded *sax-7* isoforms. Isoforms a and d, and isoforms b and c, are nearly identical, except for a short sequence of 9 extra nucleotides at the beginning of exons 17 and 14 in isoforms c and d, respectively. *sax-7* mutant alleles and primers used for RT-PCR analysis are indicated. Primer oCB990 (blue) was used to detect the long isoforms (b and c). Primer oCB991 (green) was used to detect the short isoforms (a and d). Primer oCB989§ (brown) specifically targets isoforms c and d, as its 3’ end sequence primes on the 9 extra nucleotides of isoforms c and d. Conversely, primer oCB988* (violet) specifically targets isoforms a and b, as it was designed to prime at the exon junction lacking the 9 extra nucleotides. To the right, detection of *sax-7* transcripts by RT-PCR. All RT-PCR products were confirmed by sequencing, and correspond to the predicted *sax-7* sequences. In <u>mutant</u>*<u>nj48</u>*, transcripts are detected. The *sax-7* long isoforms (b and c) RT-PCR product amplified with the primers oCB990/oCB987 is shorter than in the wild type, in accordance with the *nj48* deletion where exon 5 of *sax-7L* is deleted. As expected, (primer oCB991 falls within the *nj48* deletion), no transcript for short isoforms (a and d) were detected. In <u>mutants</u>*<u>nj53</u>*<u>and</u>*<u>eq2</u>*, the 5’ UTR and exon 1 of the *sax-7* long isoforms (b and c) are deleted, and for *<u>eq2</u>*, part of exon 2 is deleted as well. Yet, *sax-7* long (b and c) transcripts are detected in *nj53* and *eq2*. Finally, in <u>mutants</u>*<u>tm1448</u>*<u>and</u>*<u>eq1</u>*, both long (b and c) and short (a and d) transcripts are detected. Y45F10D.4 is a housekeeping gene used as an RT-PCR control (Hoogewijs et al., 2008). **(D)** Western blot analyses of SAX-7 protein. An antibody specific to the intracellular domain (ICD) of SAX-7 was used, which detects a region in the C-terminus of SAX-7S and SAX-7L (Chen et al., 2001). anti-HSP90 was used as a loading control. “+” indicates wild-type strain. Representative membranes of at least 3 independent repeats each. Asterisks (*) denote *non-specific* bands, which are unrelated to SAX-7 as they are present in extracts of: (1) *sax-7(qv30)* null mutants, where the entire *sax-7* genomic locus is deleted, and (2) *eq1* mutant, where the entire the region coding for the epitope targeted by this antibody is deleted. “?” indicates an unknown form of SAX-7, which is detected in both wild type and *sax-7* mutants, except for the null *qv30* and the epitope-control *eq1*. <u>Left panel</u>: >5000 mixed-stages worm populations were loaded per well. The band corresponding to <u>full-length SAX-7L (∼190 kDa)</u> is indicated by a blue arrow; SAX-7L is detected in the wild type and in mutants *qv25* and *qv26*, but not in *eq2* and *nj53*, as expected. The band corresponding to <u>full-length SAX-7S (∼150 kDa)</u> is indicated by a green arrow; SAX-7S is detected in the wild type and in mutants *eq2* and *nj53*, but not in *qv25* and *qv26*, as expected. A presumably truncated mutant version of SAX-7L is detected in *eq2* (blue arrowhead), which is not detected in wild type or *eq1* and *qv30* controls. Also, a truncated mutant version of SAX-7 is detected in *nj48* (black arrowhead, unclear whether it corresponds to a truncated SAX-7S and/or SAX-7L). <u>Two cleavage products</u> are also detected: a highly abundant band at ∼60 kDa, indicated by a red arrow, corresponds to the C-terminal product resulting from cleavage at the serine protease site; a less abundant band at ∼28 kDa, indicated by a black arrow, corresponds to the C-terminal product from the cleavage site near to the transmembrane domain, and appears to run as a double band. An exposure of 9.5 sec is required to see the bands of full-length SAX-7L and SAX-7S (arrows); however, at this exposure, the ∼60 kDa cleavage product saturates the area indicated by the red dotted box. To better distinguish level differences among mutants, the same ∼60 kDa membrane area was exposed for 0.1 sec and is shown underneath. In SAX-7S-specific mutants *qv25* and *qv26*, the 60 kDa C-terminal product (resulting from cleavage at the serine protease site) is detected at a lower level of compared to wild type; however, in SAX-7L-specific mutants *eq2* and *nj53* the level of this 60 kDa C-terminal serine protease cleavage products is comparable to wild type, suggesting that most of this cleavage product may be derived from full-length SAX-7S. On the other hand, the 28 kDa C-terminal cleavage (resulting product from cleavage site near to TM) appears to be less abundant in *eq2*. <u>Right panel</u>: 100 L4 worms (4^th^ larval stage) were loaded per well. While not all protein forms can be detected with this lower protein amount, the 60 kDa C-terminal product from cleavage at the serine protease site is again clearly detected at lower levels in the *sax-7S*-specific mutants *qv25* and *qv26*, compared wild type.

We also carried out western blots to characterize the expression of the protein SAX-7 in *sax-7(nj48)* and other mutant alleles (**Fig. 1D**). To detect SAX-7, we used a purified antibody generated against the cytoplasmic tail of SAX-7 (Chen et al., 2001). In wild-type extracts, we detect five protein bands of ∼190 kDa, 150 kDa, 60 kDa, 40 kDa and 28 kDa that are absent in the control *eq1*, in which the epitope-containing region of SAX-7 is deleted, and in a newly generated deletion allele *qv30*, in which the entire locus of *sax-7* is absent (see below). The 190 kDa band (**Fig. 1D**, blue arrow) and the 150 kDa band (**Fig. 1D**, green arrow) correspond to the predicted SAX-7L and SAX-7S full-length protein, respectively, as previously reported (Chen et al., 2001; Sasakura et al., 2005; Wang et al., 2005). Two bands at 60 kDa and 28 kDa appear to be cleavage products. The 60 kDa band (**Fig. 1D**, red arrow) is likely the C-terminal fragment resulting from proteolytic cleavage of SAX-7 at the serine protease site in the 3^rd^ FnIII domain (**Fig. 1B**). This cleavage site is conserved in vertebrate L1 proteins (Faissner et al., 1985; Haspel and Grumet, 2003; Hortsch, 1996, 2000; Kalus et al., 2003; Lutz et al., 2017; Lutz et al., 2012; Matsumoto-Miyai et al., 2003; Mechtersheimer et al., 2001; Nayeem et al., 1999; Sadoul et al., 1988; Schafer and Altevogt, 2010; Silletti et al., 2000; Xu et al., 2003). The 28 kDa band (**Fig. 1D**, black arrow), which runs as a doublet, is likely the predicted C-terminal fragment resulting from the proteolytic cleavage of SAX-7 at the proximal-transmembrane extracellular site (TM site, **Fig. 1B**). Similar metalloprotease cleavage sites have been reported in vertebrate L1CAM proteins (Beer et al., 1999; Gutwein et al., 2003; Haspel and Grumet, 2003; Jafari et al., 2010; Kalus et al., 2003; Kiefel et al., 2012; Linneberg et al., 2019; Maretzky et al., 2005; Maten et al., 2019; Matsumoto-Miyai et al., 2003; Mechtersheimer et al., 2001; Naus et al., 2004; Nayeem et al., 1999; Riedle et al., 2009; Sadoul et al., 1988; Schafer and Altevogt, 2010; Tatti et al., 2015; Xu et al., 2003; Zhou et al., 2012). Finally, a 40 kDa band is detected in the wild type, but is absent in the controls (**Fig. 1D**), suggesting yet another form of SAX-7 (also recently indicated in WormBase). Noteworthy, we find that the level of the full-length SAX-7L protein is higher than the full-length SAX-7S, and that most SAX-7 is detected as a cleaved form. In particular, the serine protease-cleavage product (∼80%) is most abundant (only the C-terminal fragment of the serine protease cleavage can be detected, as epitope located in the C-terminus). In contrast, the proximal-TM cleavage site product is less abundant (**Fig. 1D**). Importantly, in extracts of *nj48* mutants, two forms of SAX-7 protein were detected, which were absent in controls: a 40 kDa band (**Fig. 1D**, indicated by a question mark), and a 140 kDa band, likely a truncated form of SAX-7 protein (**Fig. 1D**, black arrowhead). Thus, *sax-7* transcript and SAX-7 protein are detected in extracts of *sax-7(nj48)* mutants, revealing that *nj48* is not a null allele.

**Table 1.** List of *sax-7* mutant alleles used.

| Allele | Nature of alleles | Location on<br>cosmid C18F3 | Reference |
| --- | --- | --- | --- |
| **qv31** | 732 bp insertion<br>sfGFP::SAX-7S construct | after 12809 | This study |
| **qv30** | 19,972 bp deletion<br>Total loss of function | 8364-28335 | This study |
| **qv25** | 47 bp insertion, which<br>creates an ORF frameshift and a stop<br>codon in *sax-7S* signal peptide | after 12785 | This study |
| **qv26** | 36 bp insertion<br>In frame but disrupts *sax-7S* signal<br>peptide | after 12785 | This study |
| **eq2** | 648 bp deletion | 8041-8688 | (Wang et al., 2005) |
| **nj53** | 724 bp deletion | 8122-8845 | (Sasakura et al., 2005) |
| **nj48** | 582 bp deletion | 12457-13038 | (Sasakura et al., 2005) |
| **tm1448** | 1,727 bp deletion | 22599-24325 | Mitani lab at NBRP<br>*C. elegans* |
| **eq1** | 2,020 bp deletion | 26591-28605 | (Wang et al., 2005) |

### Generation of a *sax-7* null and *sax-7S*-specific mutant alleles

To generate a null allele of *sax-*7, we used CRISPR-Cas9 technology and deleted the entire locus of the *sax-7* gene. Two targets were used, one on the 1^st^ exon of *sax-7L* and one on the last exon of *sax-7* (exon 17 and 14 of the long and short isoform, respectively), resulting a 19,972 bp deletion (**Figs. 1A, S1A**). This new mutant, named *sax-7(qv30)*, is a clear null allele of *sax-7* and was verified by multiple PCRs, sequencing, and western blot (**Fig. 1D**). *sax-7(qv30)* null mutants are viable and have a somewhat reduced brood size, but their egg laying and embryonic viability are normal (**Fig. S2**).

We also generated *sax-7S*-isoform specific alleles, as this isoform has been found to be functionally important. Using CRISPR-Cas9 technology, we targeted the 1^st^ exon of *sax-7S* specifically (in a region corresponding to an intron in *sax-7L*) and obtained two small *sax-7S*-specific insertion alleles, *qv25* and *qv26*, both predicted to be strong loss-of-function alleles of *sax-7S. qv25* has a 47 bp insertion and qv26, a 36 bp insertion (**Fig. S1B-C**). Both alleles disrupt the *sax-7S* export signal peptide sequence, likely disturbing SAX-7S protein synthesis. As a further consequence of the *qv25* insertion, a stop codon is generated in the open reading frame of *sax-7S* (**Fig. S1B**). At the protein level, using the antibody against the SAX-7 cytoplasmic tail (Chen et al., 2001), as expected we detected no full-length SAX-7S in extracts of these mutants, while full-length SAX-7L was detected (190 kDa band; **Fig. 1D**). As a note, it appears that when SAX-7S is affected, as in *qv25* and *qv26*, the 60 kDa-C-terminal-serine protease-cleavage product is less abundant than in wild type or *sax-7L*-specific mutants *eq2* and *nj53* (60 kDa band; **Fig. 1D**). This was consistently observed in all of the western blots done using either mixed worm populations or 100 L4 worms (≥3 independent repeats in each case). It thus appears that a large proportion of the C-terminal serine protease cleavage product may originate from cleavage of SAX-7S protein specifically.

### Phenotypic characterization of new *sax-7* mutants

We characterized the phenotypic consequences of the complete loss of *sax-7* function in *sax-7(qv30)* mutants in neuronal maintenance. As a measure of head ganglia organization, we examined two pairs of head chemosensory neurons (ASH and ASI) from the 2^nd^ larval stage to adulthood, as previously done for other mutants (Benard et al., 2009; Benard et al., 2012; Benard et al., 2006). The soma of these neurons are located in the lateral head ganglia and their axons project into the nerve ring. We visualized these 4 neurons using the fluorescent P*sra-6::gfp* or P*sra-6::DsRed2* and noted the relative position of the ASH/ASI cell bodies with respect to the nerve ring. We found that head ganglia organization is normal in 2^nd^ larval stage *qv30* null mutants, but that it becomes progressively disorganized by the 4^th^ larval stage, worsening into adulthood (**Fig. 2A**). Similar disorganization of ASH/ASI has been described in *nj48* mutant adults (Benard et al., 2012).

**Fig 2.**
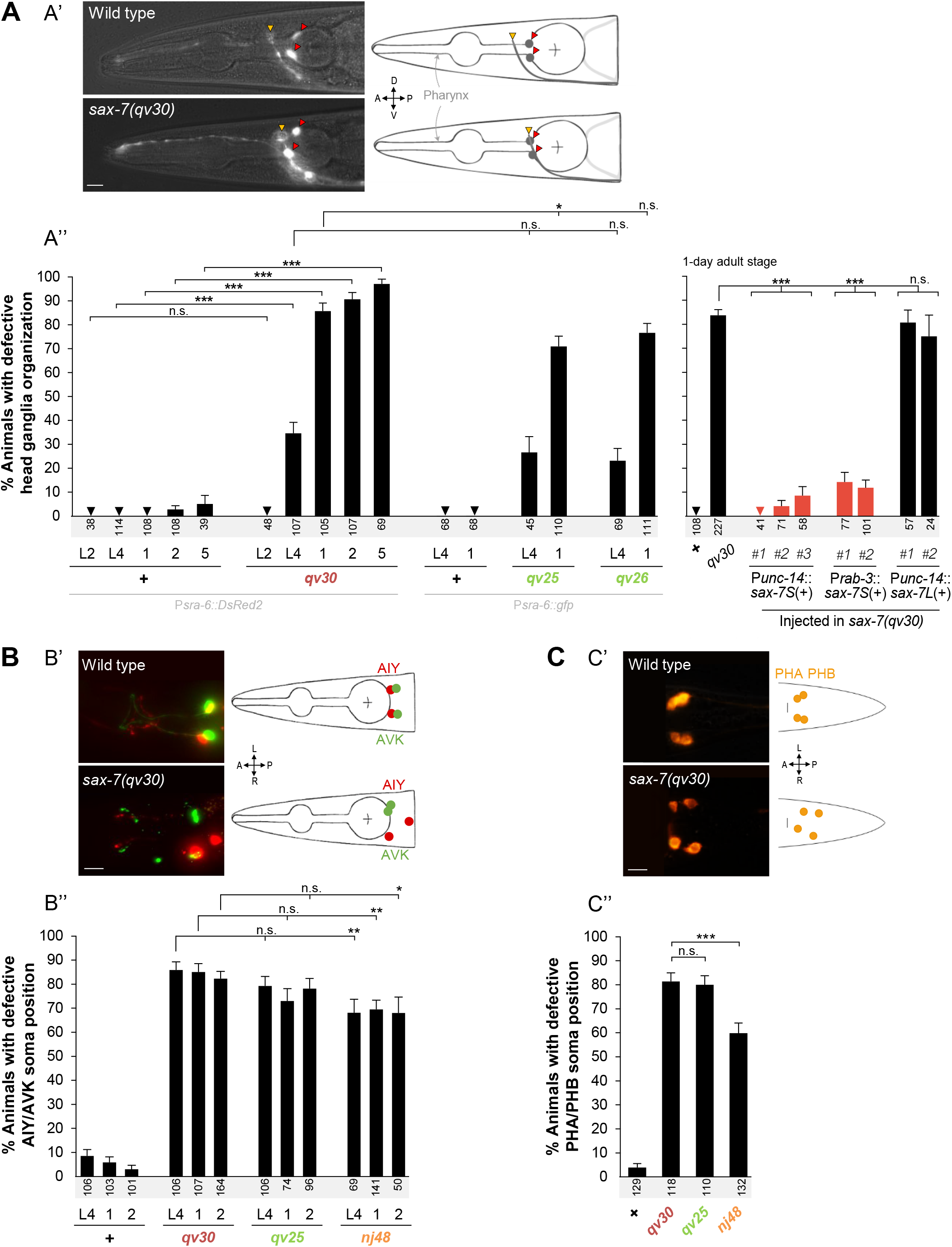
Neuronal maintenance defects in the *sax-7* mutant alleles *qv30, qv25* and *qv26*. **(A)** *sax-7S* is required to maintain head ganglia organization post-developmentally. **(A’)** Fluorescence images of the head region, where the soma and axons of the chemosensory neurons ASH and ASI are visualized using reporter P*sra-6::DsRed2*. Drawings illustrate microscopy images. Reporters P*sra-6::DsRed2* (*hdIs29*) and P*sra-6::gfp* (*oyIs14*) give comparable results for all genotypes tested. In the wild type, the soma of neurons ASH/ASI (red arrowheads) are positioned posteriorly relative to the nerve ring (yellow arrowhead) throughout stages. In *sax-7* mutants, the relative positioning between the soma of neurons ASH/ASI and the nerve ring is initially normal (soma posterior to nerve ring), but becomes progressively defective in late larvae onwards (soma can either overlap with or become anterior to the nerve ring). (**A’’**) Quantification of the relative positioning between the ASH/ASI soma and the nerve ring in wild type, null mutant *qv30*, and *sax-7S*-specific mutants *qv25* and *qv26*. Animals were examined at the 2^nd^ (L2) and 4^th^ (L4) larval stages, as well as at days 1, 2, or 5 of adulthood. Rescue of *qv30* null mutant defects by expression of *sax-7S*(+) in the nervous system using the heterologous promoters P*unc-14* and P*rab-3* (expression of *sax-7L*(+) does not rescue). Relative positioning between the soma of neurons ASH/ASI and the nerve ring was examined at 1-day adulthood using reporter P*sra-6::DsRed2*. Statistical comparisons are with *qv30* mutant. **(B)** *sax-7S* is required to maintain the retrovesicular ganglion organization. (**B’**) Fluorescence images showing the soma of two pairs of interneurons AVK and AIY on either sides of the animal, visualized using reporters P*flp-1::gfp* and P*ttx-3::DsRed2*. In the wild type animals, the soma of AVK (green) and AIY (red/yellow when overlap) are adjacent with each other. In *sax-7* mutants, one or both of the AVK and AIY neuron pairs become separate from one another. (**B’’**) Quantification of animals showing separate pairs of AVK and AIY soma in wild type, null mutant *qv30, sax-7S*-specific mutant *qv25*, and hypomorphic mutant *nj48*, at the 4^th^ larval stage (L4) and days 1 and 2 of adulthood. The *qv30* null and *qv25 sax-7S*-specific mutants are more affected than *nj48* mutants. **(C)** *sax-7S* functions to maintain tail ganglia organization. (**C’**) Fluorescence images of the chemosensory neurons PHA and PHB, visualized using DiI staining, whose soma are located in the lumbar ganglia on each side of the animal. In the wild type, the PHA and PHB soma are adjacent to each other. In *sax-7* mutants, one or both of the PHA/PHB pairs are separated from one another. (**C’’**) Quantification of disorganized soma position in wild-type, null mutant *qv30, sax-7S*-specific mutant *qv25*, and hypomorphic mutant *nj48*, at the 4^th^ larval stage. The *qv30* null and *qv25 sax-7S*-specific mutants are more severe than *nj48* mutants. Scale bar, 10 μm. Sample size is indicated under each column of the graph. Error bars are standard error of the proportion. Asterisks denote significant difference: * p ≤ 0.05, ** p ≤ 0.01, *** p ≤ 0.001. (z-tests, p values were corrected by multiplying by the number of comparisons, Bonferroni correction). “+” indicates wild-type strain; n.s., not significant.

We also examined the precise axon position of the two pairs of bilateral interneurons (PVQ and PVP) in the ventral nerve cord, labelled by the reporters P*sra-6::DsRed2* and P*odr-2::cfp*, respectively. These axons are normally positioned in freshly hatched 1^st^ stage larvae of *qv30* mutants, indicating that they had extended normally along the ventral nerve cord during embryogenesis. However, compared to wild type where the PVQ and PVP axons remain well positioned in virtually all animals (94%, n=117), in *sax-7(qv30)* mutants these axons later fail to maintain this positioning and inappropriately flip-over to the other side of the ventral nerve cord in 37.5% of *qv30* animals (n=80), which is similar to *nj48* mutants (Benard et al., 2012; Pocock et al., 2008).

Other aspects of neuroanatomy of *qv30* mutants were more severe than *nj48* mutants. For instance, we observed retrovesicular ganglia organization by visualizing the neurons AIY and AVK (using reporters P*ttx-3::mCherry* and P*flp-1::gfp*, respectively) and found that 85% of 1-day adult *qv30* mutant animals display disjointed AIY and AVK soma, compared to 70% in *nj48* mutants (**Fig. 2B**). Also, using DiI staining we found that the position of the soma of PHA and PHB in the tail ganglia is defective in 81% of *qv30* mutants, compared to 60% of *nj48* mutants, at the 4^th^ larval stage (**Fig. 2C**). Thus, while *nj48* is a strong allele displaying similar penetrance to the null allele *qv30* in some neuronal contexts, its loss of function is partial and less severe than the null *qv30* in other neuronal contexts.

### SAX-7S is required for neuronal maintenance

*sax-7S*, but not *sax-7L*, has previously been found to be sufficient to rescue neuronal maintenance defects in *sax-7(nj48)* mutants (Pocock et al., 2008; Sasakura et al., 2005). We verified whether *sax-7S* is also sufficient to rescue such defects in the *sax-7(qv30)* null mutants, by generating transgenic *qv30* null mutant animals carrying wild-type copies of *sax-7S*(+) expressed neuronally [using the transgenes P*unc-14*::*sax-7S*(+) and P*rab-3::sax-7S*(+)]. We found that *qv30* transgenic animals were profoundly rescued for head ganglia disorganization (**Fig. 2A**). On the other hand, wild-type *sax-7L*(+) did not rescue *qv30* transgenic mutant animals (transgene P*unc-14*::*sax-7L*(+); **Fig. 2A**), similar to findings using the allele *nj48* (Pocock et al., 2008; Sasakura et al., 2005). This is consistent with the absence of defects in *sax-7L*-specific mutants *eq2* and *nj53* (Benard et al., 2012; Sasakura et al., 2005). Altogether, these results further demonstrate that *sax-7S* mediates neuronal maintenance function.

To directly assess the phenotypic consequences of specifically disrupting *sax-7S*, we analyzed neuronal maintenance defects of the newly generated *sax-7S*-specific mutants *qv25* and *qv26* (**Figs. 1A, 2A**). We found that the severity of their defects is similar to *qv30* null mutant animals. For instance, the head ganglia of *qv25* and *qv26* animals become disorganized from the 4^th^ larval stage onwards, similar in penetrance and expressivity to the *qv30* null mutants (**Fig. 2A**). Also, the soma of retrovesicular ganglion neurons AIY and AVK become disorganized from the 4^th^ larval stage in *qv25* mutants, similar to *qv30* mutants (**Fig. 2B**). Finally, the soma of tail neurons PHA and PHB get displaced from the 4^th^ larval stage onwards in *qv25* mutants, similar to *qv30* mutants (**Fig. 2C**). Thus, the specific disruption of *sax-7S* leads to neuronal maintenance defects that are similar to those resulting from the complete loss of *sax-7* (deleting both *sax-7S* and *sax-7L*), confirming the key role of SAX-7S in the maintenance of neuronal architecture.

### Post-developmental expression of *sax-7S* is sufficient for maintaining neuronal architecture

Although the ventral nerve cord and head ganglia assemble during embryogenesis, *sax-7(qv30)* null mutants manifest ventral nerve cord flip-over defects during larval development, and head ganglia become disorganized by late larval stages, progressively worsening into adulthood. The appearance of defects in *sax-7* mutants could in theory result from either (a) undetected embryonic neuronal development defects that later worsen as the animal grows and moves, or (b) deficient neuronal maintenance during larval and adult stages. To distinguish between these possibilities, we carried out rescue assays of *qv30* null mutants with wild-type *sax-7S*(+) copies expressed under the control of an inducible heat shock promoter, which drives expression in neurons and other tissues (Fire et al., 1990; Jones et al., 1986; Stringham et al., 1992). For this, we generated transgenic *qv30* animals carrying the transgene P*hsp16*.*2::sax-7S*(+) as an extrachromosomal array. All animals were kept at 15°C [a colder temperature to prevent expression of P*hsp16*.*2::sax-7S*(+)], except during heat shock treatments (**Fig. 3A**). The organization of the ASI and ASH head ganglia neurons was examined in 1-, 2-, 3-, 4-, and 5-day old adults. We controlled for head ganglia organization in the strains grown continuously at 15°C being indeed (a) normal in wild-type animals; (b) defective in *qv30* mutants; (c) not rescued in the absence of heat shock, in transgenic *qv30* animals carrying the transgene [P*hsp-16*.*2::sax-7S*(+)], indicating that the transgene is not expressed without heat shock; and (d) defective in *qv30* non-transgenic control siblings under the same conditions (**Fig. 3B**).

**Fig 3.**
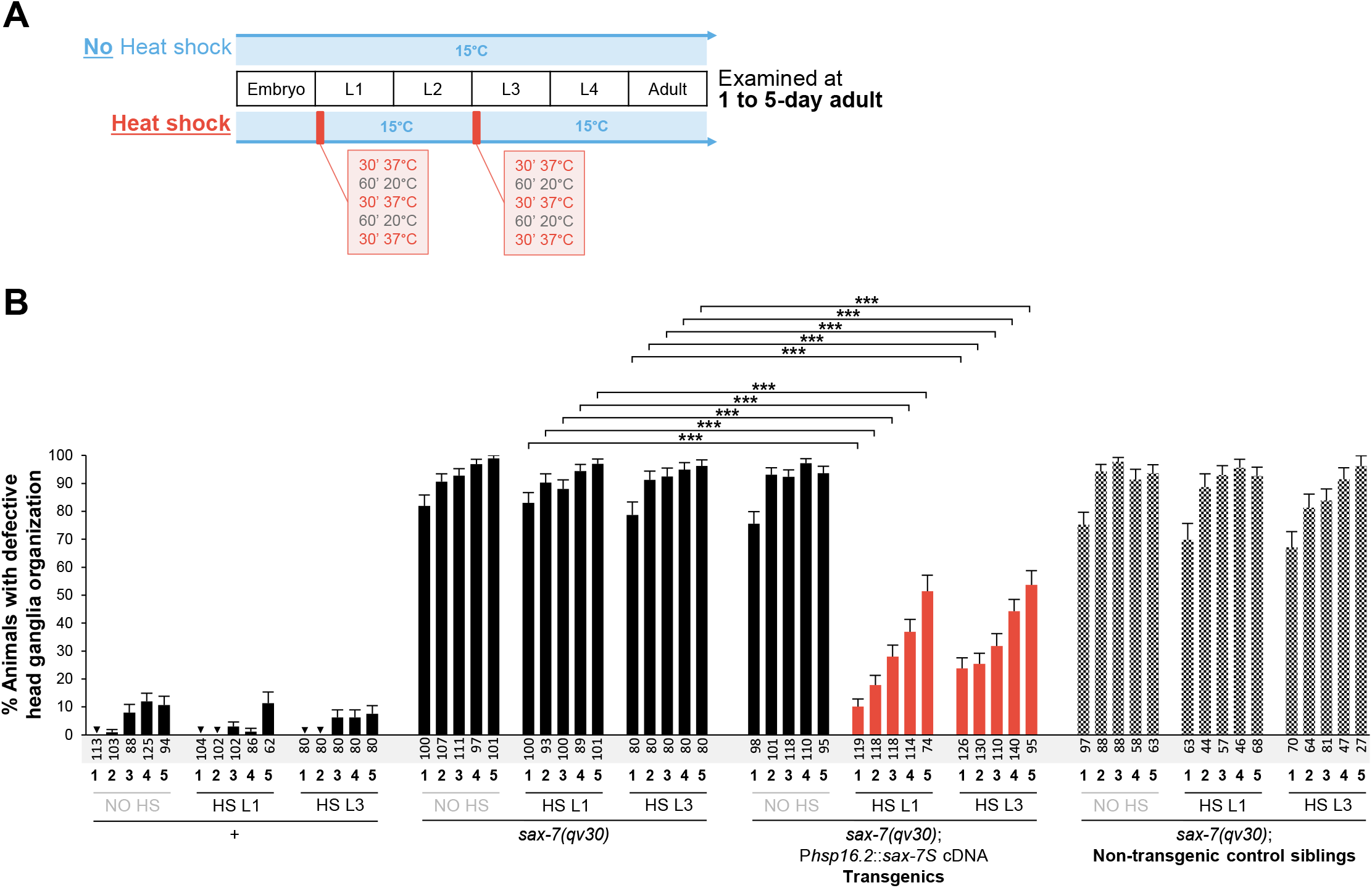
Expression of *sax-7S*(+) during larval stages is sufficient for *sax-7S* to function in the maintenance of neuronal organization. **(A)** Summary of the heat shock experiments performed. Animals were kept at 15°C at all times except during heat shock at 37°C (red boxes). Heat shock was done at either the 1^st^ (L1) or the 3^rd^ (L3) larval stage. Animals were later analyzed at days 1, 2, 3, 4, and 5 of adulthood. **(B)** Quantification of the relative position between the soma of ASH/ASI and the nerve ring (as in Fig. 2A), visualized using the reporter *oyIs14* (P*sra-6::gfp*), at days 1, 2, 3, 4 and 5 of adulthood (age indicated under each bar of the graph). Transgenic animals carry a transgene of *sax-7S*(+) expressed under the control of a heat-shock promoter (P*hsp-16*.*2::sax-7S*(+)). Controls include the wild type, *sax-7(qv30)* mutants, and non-transgenic siblings of the transgenic animals, which are derived from the same mothers and grew on the same plates, but which do not carry the extrachromosomal array harboring the transgene. Additionally, for all of the four genetic conditions, neuroanatomical analyses were done in the absence of heat shock so as to ensure that no transgene expression occurred in the absence of heat shock. The defects that adult *sax-7(qv30)* mutants normally display are profoundly rescued by heat-shock-induced expression of *sax-7S*(+) at larval stages, as seen in heat-shocked adult *sax-7(qv30)* mutants carrying the transgene, (orange bars). Non-transgenic siblings, however, are severely defective, indicating that the rescue of defects is dependent on expression of *sax-7S(+)* upon heat shock. “+”, indicates wild-type; “NO HS”, no heat-shock; “HS L1”, heat shock was performed at the 1^st^ larval stage; “HS L3”, heat shock was performed at the 3^rd^ larval stage. Sample sizes is indicated along the grey zone, under each bar of the graph. Error bars are standard error of the proportion. Asterisks denote significant difference: *** p ≤ 0.001. (z-tests, p values were corrected by multiplying by the number of comparisons, Bonferroni correction).

To determine the temporal requirement in *sax-7* function, we heat shocked 1^st^ (L1) or 3^rd^ (L3) larval stage animals that had otherwise been grown at 15°C, and examined head ganglia organization at days 1, 2, 3, 4, and 5 of adulthood (**Fig. 3A**). Wild-type animals, *qv30* mutants, transgenic *qv30* animals carrying the transgene [P*hsp-16*.*2::sax-7S*(+)], and their *qv30* non-transgenic siblings, were analyzed in parallel. An additional control consisted of heat-shock treatment alone, in the absence of the transgene, which did not modify the defects of *sax-7* mutants (head ganglia are similarly disorganized in *qv30* animals whether heat shocked or not; **Fig. 3B**). Also, heat shock did not alter head ganglia organization in wild-type animals (**Fig. 3B**). In contrast, when transgenic *qv30* animals carrying the transgene P*hsp16*.*2::sax-7S*(+) were heat shocked at L1, or even as late as L3, their neuronal organization was profoundly rescued. This rescue by heat shock-induced expression of *sax-7S(+)* is dependent on the presence of the transgene, as non-transgenic control siblings (which grew side by side with *qv30* transgenics) were not rescued (**Fig. 3B**). Together, these results indicate that wild-type activity of *sax-7S* provided as late as the 3^rd^ larval stage is sufficient for it to function in the maintenance of neuronal architecture. Thus, *sax-7S* can function post-developmentally to maintain the organization of embryonically developed neuronal architecture. Moreover, we found that the rescue of *qv30* mutants following induction of *sax-7S*(+) is more profound in younger adults (days 1 to 3), as compared to older adults (days 4 and 5, **Fig. 3B**). By day 5 of adulthood, more than 6 days have passed after heat shock-induced expression of P*hsp16*.*2::sax-7S*(+), suggesting that *de novo* expression of *sax-7S* may be required to ensure its maintenance function during adulthood.

### Endogenous *sax-7S* is expressed in neurons

The *sax-7* gene is expressed strongly and broadly across the nervous system, as visualized with a fosmid (Sarov et al., 2012) where both the short and long isoforms are tagged with *gfp* (**Fig. 4A**; (Ramirez-Suarez et al., 2019)). To elucidate the expression pattern of *sax-7S*, we used CRISPR-Cas9 technology to tag the *sax-7S* isoform specifically with *sfgfp* at its endogenous genomic locus (**Fig. 4B**). We targeted the end of the 1^st^ exon of *sax-7S* in a precise region that corresponds to intron 4 of *sax-7L*, and inserted *sfgfp*, preceded by *sax-7S*-signal-peptide coding sequence (sfGFP::SAX-7S; **Figs. S1D, 4B**). This knock-in allele of *sax-7*, named *qv31*, which was verified by sequencing (**Fig. S1D**), does not affect overall morphology or behavior, and head ganglia organization of *qv31* animals is normal (n=84, 2% defects, examined at 1-day adult by DiI staining, similar to wild type with 0% defects, n=76).

**Fig 4.**
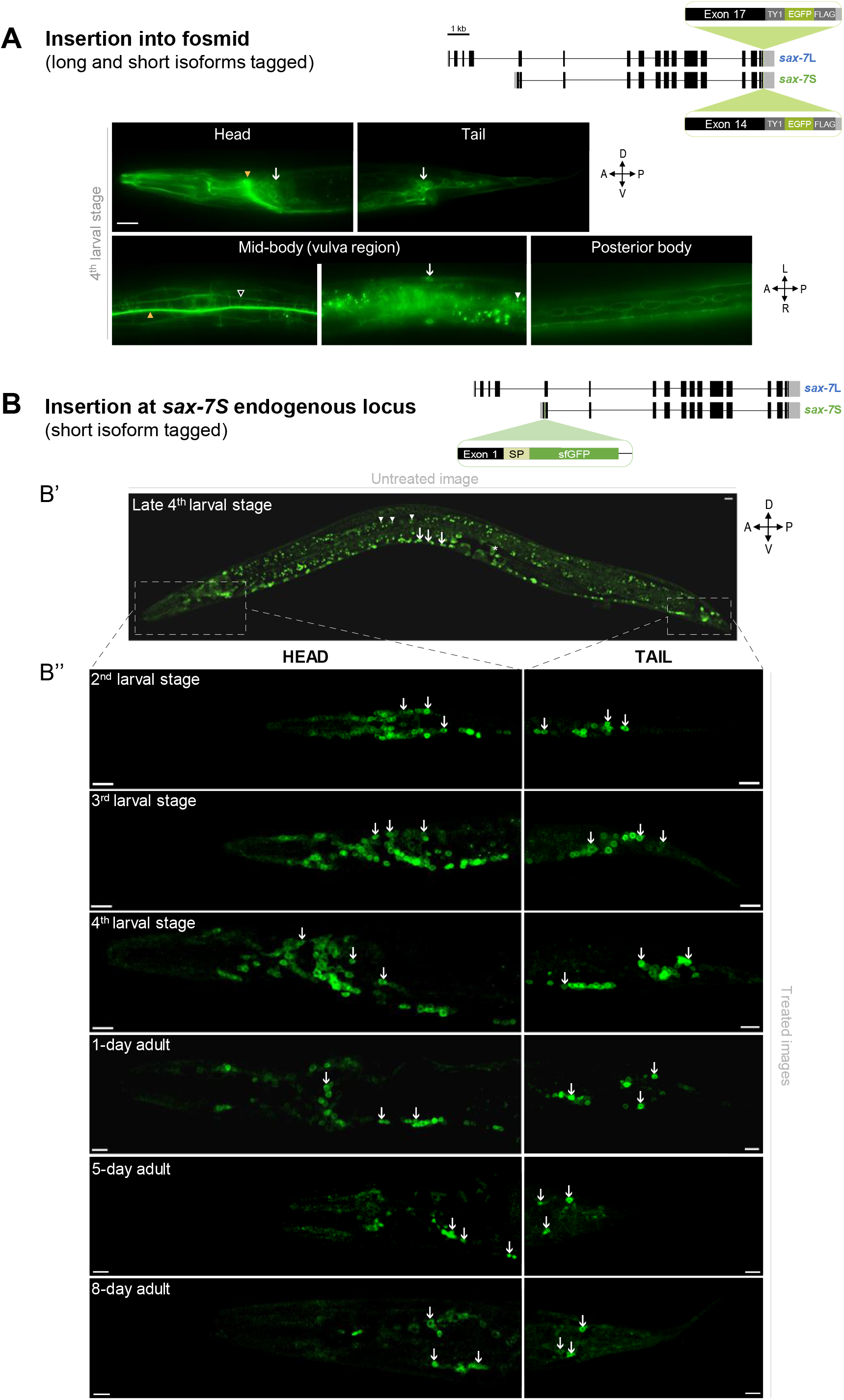
SAX-7S is expressed in virtually all neurons throughout life. **(A)** Images of SAX-7::GFP expression reporting both SAX-7L and SAX-7S. As shown on the schematics, in this previously published transgene (Sarov et al., 2012), the gene coding for EGFP was inserted into the gene *sax-7* by fosmid recombineering in such a way that both SAX-7S and SAX-7L isoforms were tagged, making impossible to distinguish between them. SAX-7::GFP is broadly expressed in neurons and epidermal cells (vulval cells, seam cells). **(B)** Confocal images showing sfGFP::SAX-7S expression. As shown on the schematics, the gene coding for sfGFP was inserted by CRISPR-Cas9 at the end of exon 1 of *sax-7S* in order to specifically tag SAX-7S (see **Fig. S1D**; *qv31* in **Table 1**). “sfGFP”, superfolderGFP; “SP”, export signal peptide sequence part of *sax-7S* inserted along with *sfgfp*. (**B’**) Untreated confocal image of a late 4^th^ larval stage worm. Arrows indicate neurons of ventral nerve cord and arrowheads point to examples of background green auto-florescence due to gut granules. Dotted boxes indicate the body region (head or tail) analyzed in B”. (**B”**) Images of animals at the indicated larval stages and days of adulthood, examined by confocal microscopy followed by unmixing. Aged worms (>5-days old) have notably increased background auto-fluorescence. Arrows indicate sfGFP::SAX-7S expression in neurons of the head (left) or tail (right) ganglia. n ≥ 20 animals examined by confocal microscopy for each stage. z-stack projections. Scale bar, 10 μm.

To characterize the temporal and spatial expression pattern of *qv31* sfGFP::SAX-7S, we used conventional as well as confocal fluorescence microscopy with spectral unmixing. sfGFP::SAX-7S is seen most predominantly and abundantly across the nervous system, where it is observed in virtually all neuron of head and tail ganglia, the ventral nerve cord, as well as in isolated neurons located along the body wall (e.g. HSN near the vulva, and the PVM post-deirid neuron). Expression of sfGFP::SAX-7S in neurons is first observed in embryos (**Fig. S3A**), and persists throughout larval stages (**Fig. 4B**) and adulthood, including in 5- and 8-day adults (**Fig. 4B**). While virtually all neurons express SAX-7S, differences in the level of expression are observed among neurons. sfGFP::SAX-7S is also occasionally detected in other cell types, such as in epidermal cells of the developing vulva and the uterus at the L4 stage, but not in adults (**Fig. S3B**). In sum, SAX-7S appears to be transiently and weakly expressed in developing cells of the epidermis, but its expression is strongest and sustained in virtually all neurons from embryogenesis to adulthood.

As a note, previous reports where *sax-7* (both L and S indistinctly) was tagged intracellularly reported SAX-7 protein signal in axons, dendrites or the plasma membrane (Chen et al., 2001; Ramirez-Suarez et al., 2019; Wang et al., 2005). Here, the sfGFP::SAX-7S signal in *qv31* animals appears to be peri-nuclear in neuronal cell bodies, which is surprising for a transmembrane protein, and is likely artifactual. Indeed, in our effort to exclusively tag SAX-7S with sfGFP by CRISPR-Cas9, the only option was to insert the sfGFP very close to the predicted signal peptide of SAX-7S, which possibly affects the cleavage of the signal peptide or targeting of the protein. Thus, *qv31* does not reliably inform about the subcellular localization of the protein SAX-7S, yet it yields valuable information about the spatio-temporal expression pattern of *sax-7S*.

### Domains Ig3-4 of SAX-7S are necessary for its function in neuronal maintenance

L1 family members play diverse roles via homophilic interactions through their extracellular domains which leads to homophilic cell adhesion (Brummendorf et al., 1998; Brummendorf and Rathjen, 1996; Haspel and Grumet, 2003; Hortsch, 2000), and mutating different extracellular Ig-like domains of vertebrate L1 perturbs its homophilic and/or heterophilic binding in *in vitro* assays (Blaess et al., 1998; Castellani et al., 2002; De Angelis et al., 1999; De Angelis et al., 2002; Felding-Habermann et al., 1997; Haspel et al., 2000; Holm et al., 1995; Kunz et al., 1998; Montgomery et al., 1996; Oleszewski et al., 1999; Zhao and Siu, 1995) and neurite outgrowth (Appel et al., 1993). In *C. elegans*, neuronal expression of a SAX*-*7S recombinant version lacking Ig5-6 domains rescued AIY/AVK neuronal soma position defects of *sax-7(nj48)* mutants, whereas a recombinant version lacking Ig3-4 domains does not (Pocock et al., 2008). We asked whether such SAX-7S recombinant versions lacking specific Ig domains could rescue head ganglia organization in *qv30* null mutant animals. We found that the transgene P*unc-14::sax-7S*ΔIg5-6 rescued the position of the soma of the neurons ASH and ASI relative to the nerve ring in *qv30* null mutants, but that the transgene P*unc-14::sax-7S*ΔIg3-4 did not rescue (**Fig. 5C**). This indicates that only Ig domains 3 and 4 of SAX-7S are required for its role in the maintenance head ganglia organization.

**Fig 5.**
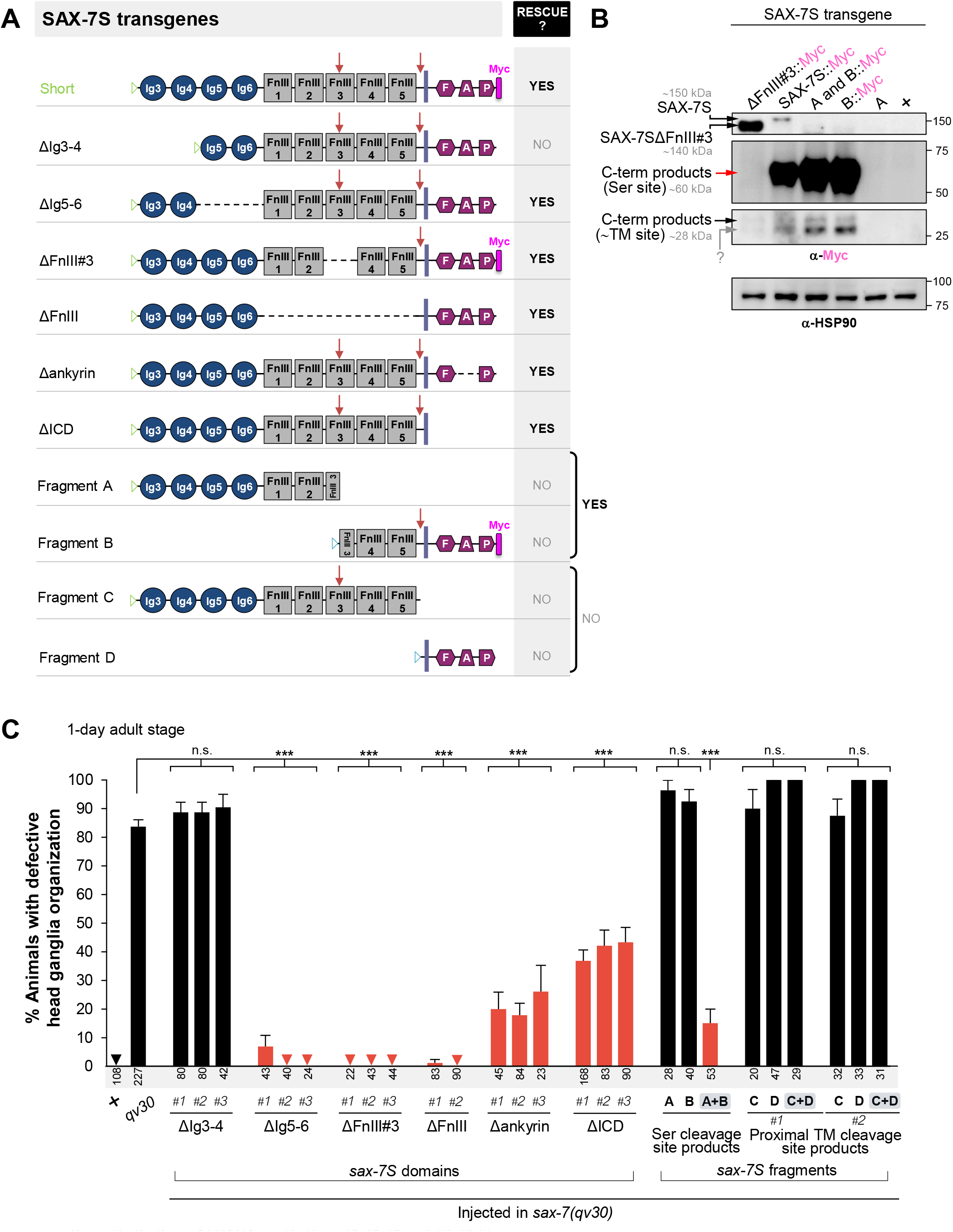
The two SAX-7S cleavage products derived from the serine protease cleavage site, together, can mediate the maintenance of neuronal architecture. **(A)** Schematics of full-length and recombinant transgenic versions of SAX-7S used in this study. Blue triangles indicate the signal peptide of SAX-7L. Green triangles indicate the signal peptide of SAX-7S. “<u>ΔIg3-4</u>” contains the entire SAX-7S protein except for the two first Ig domains. “<u>ΔIg5-6</u>” contains the entire SAX-7S protein except for the Ig5 and 6 domains. In “<u>ΔFnIII#3</u>”, SAX-7S::Myc lacks the 3^rd^ FnIII domain. In “<u>ΔFnIII</u>”, SAX-7S lacks all FnIII domains. In “<u>Δankyrin</u>”, SAX-7S lacks the intracellular ankyrin binding domain. In “<u>ΔICD</u>”, SAX-7S lacks the intracellular domain. “<u>Fragment A</u>” contains the SAX-7S protein region from Ig3 to the serine protease cleavage site (RWKR). “<u>Fragment B</u>” contains the SAX-7S::Myc protein region from the serine protease cleavage site (RWKR) to PDZ::Myc. “<u>Fragment C</u>” contains the SAX-7S protein region from Ig3 to the proximal-transmembrane cleavage site. “<u>Fragment D</u>” contains the SAX-7S protein region from the proximal-transmembrane cleavage site to PDZ. “Ig”, Immunoglobulin-like domain; “FnIII”, Fibronectin type III domain; “F”, FERM domain binding motif; “A”, Ankyrin binding motif; “P”, PDZ domain binding motif; bold violet line indicates the transmembrane domain; red arrows indicate serine protease cleavage site in FnIII#3 or, cleavage site close to the transmembrane domain. **(B)** Western blot analysis of wild-type animals (+), *sax-7(qv30)* null mutants expressing transgenes for various Myc-tagged SAX-7S fragments. N-terminal and C-terminal fragments of SAX-7S proteins were detected with anti-Myc antibody. Mixed-stage populations of >5000 worms were loaded per well, including a variable proportion of animals that actually carry the extrachromosomal array (and therefore are transgenic), as the array gets lost randomly upon cell divisions and generations; this comparison is only qualitative. As expected, in lysates of worms with transgene SAX-7SΔFnIII#3, an uncleaved band smaller than the full-length SAX-7S is detected. “C-term products” indicates C-terminal cleavage product, “Ser site” indicates serine protease cleavage site, “βTM site” indicates cleavage site near to transmembrane domain. “?” indicates an unknown form of SAX-7S. The three top anti-Myc panels correspond to the same membrane but at different exposure times in order to facilitate the observation of bands that are largely different in abundance (as was done in Fig. 1D). α-HSP90 was used as a loading control. **(C)** The defects of *qv30* null mutants are rescued by the expression of specific *sax-7S*(+) variants in the nervous system using the heterologous promoter P*unc-14*. The relative positioning of the soma and nerve ring axons of chemosensory neurons ASH/ASI (as in Fig. 2A) was evaluated using the reporter P*sra-6::DsRed2*. Wild-type control and *qv30* mutants, along with distinct SAX-7S recombinant transgenic animals, were examined as 1-day adults. Domain analyses are shown on the left of the graph, and fragment analyses on the right, as indicated. The simultaneous absence of Ig3 and 4 fails to rescue, while other domain deletions remain fully or largely functional. For fragment analyses, fragment A and B rescue the defects of the null mutant, indicating that the two SAX-7S protein fragments somehow reconstitute function. Two separate sets of independent extrachromosomal arrays for fragments C and D were tested (C#1+D#1, and C#2+D#2), which failed to rescue. Sample size is indicated under each column of the graph. Error bars are standard error of the proportion. Asterisks denote significant difference: *** p ≤ 0.001. (z-tests, p values were corrected by multiplying by the number of comparisons, Bonferroni correction). “+”, indicates wild-type strain; n.s., not significant.

FnIII domains of L1 family members play diverse roles in neurite outgrowth, homophilic binding, and interactions with various partners (Haspel and Grumet, 2003; Holm et al., 1995; Kalus et al., 2003; Koticha et al., 2005; Maten et al., 2019; Silletti et al., 2000). We asked whether FnIII domains are necessary for *sax-7S* function in *C. elegans* to maintain head ganglia organization. The recombinant transgene P*unc-14::sax-7S*ΔFnIII#3::Myc lacks the third FnIII (FnIII#3) domain, which harbors the serine protease cleavage site (**Fig. 5A**, ΔFnIII#3). In extracts of *qv30* transgenic animals carrying this transgene, a ∼140 kDa band is detected with anti-Myc antibodies (**Fig. 5B**), which is the expected size for uncleaved SAX-7S minus the FnIII#3 domain (full-length SAX-7S would be ∼150 kDa). We found that this transgene rescues head ganglia organization defects of *qv30* mutant animals (**Fig. 5C**), indicating that uncleaved SAX-7SΔFnIII#3 can function in neuronal maintenance, at least in a transgenic overexpression situation. We next tested a transgene which lacks all five FnIII domains, P*unc-14::sax-7S*ΔFnIII, and found that it also rescues the head ganglia organization defects of *qv30* null mutants (**Fig. 5C**). Such transgene lacking all FnIII domains could also rescue AIY and AVK soma position in *nj48* mutants (Pocock et al., 2008), as well as AIY position and branching in *sax-7(dz156)* mutants (Diaz-Balzac et al., 2015).

The intracellular region (ICD) of SAX-7/L1CAM shows a strong homology between vertebrates and invertebrates, and mutations in the cytoplasmic domain leads to X-linked hydrocephalus in humans (Wong et al., 1995b). This intracellular part contains motifs (FERM, ankyrin and PDZ binding-domain motifs; **Fig. 1B**), which mediate interactions with intracellular components and cytoskeletal proteins (Davey et al., 2005; Davis and Bennett, 1994; Dirks et al., 2006; Falk et al., 2004; Gil et al., 2003; Gunn-Moore et al., 2006; Herron et al., 2009; Koroll et al., 2001; Schaefer et al., 2002; Wong et al., 1995a). We tested whether a transgene of *sax-7S* lacking the ankyrin-binding motif (P*unc-14::sax-7S*Δankyrin) could function to maintain head ganglia organization, and found that *qv30* null mutants were significantly rescued by this transgene (**Fig. 5C**). Thus, the ankyrin-binding motif does not appear to be necessary for SAX-7S function in maintenance of head ganglia. We next asked if SAX-7S could function in neuronal maintenance without its intracellular domain (P*unc-14::sax-7S*ΔICD), and found a partial but significant rescue of the *qv30* defects in head ganglia organization in animals neuronally expressing this transgene (**Fig. 5C**). However, 40% of the *qv30* animals display maintenance defects. This profound but incomplete rescue may be due to either mosaicism or overexpression of the plasmid, which could possibly interfere with interactions.

### Serine protease SAX-7S fragments can, together, function in neuronal maintenance

The most abundant detected form of SAX-7 appears to be a serine protease-cleavage product (**Fig. 1D**), which splits the molecule within the third FnIII domain (**Fig. 1B**). Other detected cleavage products result from a cleavage site proximal to the transmembrane domain (TM) (**Fig. 1D**). We tested whether the protein fragments predicted to result from serine protease-site cleavage, or the TM-proximal cleavage, could function in maintaining neuronal organization, similarly to full-length SAX-7S. For this, we constructed four separate transgenes encoding each of the four predicted fragments of the protein SAX-7S, from cleavage at either the serine protease or the TM proximal sites. Some of these transgenes encode versions of SAX-7S with a C-terminal Myc tag, which can then be examined by immunoblots (**Fig. 5A-B**). Cleavage of SAX-7S at the serine protease site within the third FnIII domain results in N- and C-terminal fragments, which we named “SAX-7S-fragment-A”, and “SAX-7S-fragment-B”, respectively (**Fig. 5A**). Another cleavage event proximal to the transmembrane domain results in N- and C-terminal fragments which we named “SAX-7S-fragment-C”, and “SAX-7S-fragment-D”, respectively (**Fig. 5A**). We tested each of these fragments alone, or in reciprocal combinations, for their ability to rescue the neuronal maintenance defects of *sax-7(qv30)* mutants when expressed under the neuronal promoter P*unc-14*. Neither “SAX-7S-fragment-A” alone (P*unc-14::sax-7S-*[N-terminal] Ig3 up to serine protease site), nor “SAX-7S-fragment-B alone” (P*unc-14::sax-7S-*serine protease site up to PDZ [C-terminal]), could rescue head ganglia organization defects of *qv30* mutant animals (**Fig. 5C**). Similarly, neither “SAX-7S-fragment-C” alone (P*unc-14::sax-7S-*[N-terminal] Ig3 up to TM site), nor “SAX-7S-fragment-D” alone (P*unc-14::sax-7S-*TM site up to PDZ [C-terminal]), could rescue head ganglia organization defects of *qv30* mutant animals (**Fig. 5C**). We next tested whether the serine protease cleavage N- and C-terminal SAX-7S fragments together, i.e. “SAX-7S-fragment-A” and SAX-7S-fragment-B” together, or whether the TM-proximal cleavage N- and C-terminal SAX-7S fragments together, i.e. “SAX-7S-fragment-C” and SAX-7S-fragment-D” together, could rescue neuronal maintenance defects of *sax-7(qv30)* mutant animals. To generate doubly transgenic animals harboring the two respective transgenes, we avoided simultaneously co-microinjecting the two transgenes, as DNA recombination events between two transgenes could potentially reconstitute a full-length gene. Thus, we instead used genetic crosses to generate doubly transgenic animals (harboring two independent extrachromosomal arrays). By genetic crosses between 2 different transgenic strains, we generated a doubly transgenic strain carrying the two extrachromosomal arrays for “SAX-7S-fragment-C” and “SAX-7S-fragment-D”, which resulted in a strain with animals carrying both extrachromosomal arrays, used for rescue assays. We found that the combination of SAX-7S fragments C and D did not rescue *sax-7(qv30)* mutant phenotype (**Fig. 5C**, “C+D”; two independent sets of extrachromosomal arrays were tested and failed to rescue). Thus, SAX-7S fragments C and D predicted to result from cleavage proximal to the TM domain, even when present simultaneously, cannot fulfill *sax-7S* function in neuronal maintenance.

In a similar manner, we crossed the strain carrying the extrachromosomal array that includes transgenic copies of “SAX-7S-fragment-A” with the second transgenic strain carrying the extrachromosomal array which includes transgenic copies of “SAX-7S-fragment-B”. The resulting strain has animals carrying both extrachromosomal arrays, which we tested for rescue of *qv30* head ganglia organization defects. The simultaneous presence of both fragments A and B, corresponding to the predicted products resulting from cleavage at the serine protease site, profoundly rescued the head ganglia organization in *sax-7(qv30)* mutant animals (**Fig. 5C**, “A+B”). This finding indicates that the two serine protease-cleavage products together can mediate neuronal maintenance. As an important control, we verified that the rescue observed in the doubly transgenic strain depends on the simultaneous presence of both transgenes (for fragments A and B). Indeed, each of the two re-derived in singly transgenic lines, each carrying only one of the two extrachromosomal arrays (fragment A alone: n=28, 82% defects, or fragment B alone: n=35, 91% defects; at 1-day adult), no longer rescued the *sax-7(qv30)* defects, further confirming that only when the two SAX-7S fragments A and B are present together in an animal can then mediate neuronal maintenance (**Fig. 5C**, “A” and “B”).

We analyzed protein extracts of doubly transgenic animals carrying both extrachromosomal arrays for fragments A and B, and as expected, no full-length SAX-7S is detected (**Fig. 5B**), confirming that the distinct SAX-7S fragments A and B, together, can fulfill the role of SAX-7S in neuronal maintenance. Together, our findings show that while the cleavage at the serine protease site is not absolutely necessary for SAX-7S function, at least in an over-expression situation, the two cleaved fragments A and B resulting from it, functionally complement to mediate normal SAX-7S function for the maintenance of neuronal architecture in *C. elegans*.

## DISCUSSION

After initial establishment of the nervous system, neuronal maintenance molecules function to actively preserve neuronal structural organization and integrity. One such molecule is *C. elegans* SAX-7, homologous to vertebrate L1 proteins, whose developmental roles have been studied (Dong et al., 2013; Salzberg et al., 2013), but whose roles in the long-term maintenance of nervous system organization remain unclear. Here we have generated and characterized a complete loss-of-function allele of *sax-7*, examined the endogenous expression pattern of SAX-7S, tested the temporal requirements for *sax-7S*, and assessed the function of SAX-7S cleavage products in the maintenance of neuronal architecture.

### New *sax-7* alleles: a complete null and two *sax-7S*-specific alleles

The *sax-7(nj48)* allele, previously considered to be a null, has detectable *sax-7* transcripts and proteins (**Fig. 1**). A new mutant, *sax-7(qv30)*, deletes the entire *sax-7* genomic locus, resulting in the complete loss-of-function of the gene (**Figs. 1A**,**D**). This null allele facilitates the interpretation of experiments without the caveat of potential truncated protein products present in hypomorphic alleles, which is especially important for rescue assays with transgenes encoding protein fragments.

*qv30* mutant animals display defects that are in some cases stronger than previously studied alleles, for instance in the maintenance of some ganglia organization (e.g., AIY and AVK neuron pairs in the retrovesicular ganglion; PHA and PHB neuron pairs in the tail ganglion; **Fig. 2**). *sax-7S*(+), but not *sax-7L*(+), can rescue the defects of null mutants *sax-7(qv30)* (**Fig. 2A**), supporting that *sax-7S* is the key isoform in maintenance of neuronal architecture, as previously described (Pocock et al., 2008; Sasakura et al., 2005). This is in accordance with previous reports that *sax-7L*-specific mutant alleles (*eq2* and *nj53*) do not lead to neuronal maintenance defects (Benard et al., 2012; Pocock et al., 2008), and that *sax-7L*(+) cannot rescue neuronal maintenance defects of mutants *nj48* and *ky146*, where both *sax-7* isoforms are affected (Pocock et al., 2008).

Our analysis of two new *sax-7* short-specific alleles, *qv25* and *qv26*, further supports the notion that *sax-7S* is the important isoform in neuronal maintenance. Mutant animals for each of these two *sax-7S* alleles display defects similar to the null *qv30* (**Fig. 2**). As a note, other *sax-7S*-specific alleles with different molecular lesions have recently been isolated (Chen et al., 2019; Rahe et al., 2019). Together, our new findings and previous results unequivocally establish that SAX-7S is the important isoform mediating maintenance of neuronal architecture (Benard et al., 2012; Diaz-Balzac et al., 2015; Pocock et al., 2008; Sasakura et al., 2005; Wang et al., 2005).

### Post-embryonic expression of *sax-7S* is sufficient to maintain head ganglia organization

Expression of *sax-7S*(+) during larval stages, which is well after the embryonic assembly of neuronal ganglia, is sufficient to function in maintaining ganglia organization (**Fig. 3**). Indeed, driving *sax-7S* expression (under the control of a heat shock promoter) at the 1^st^ larval stage, or as late as the 3^rd^ larval stage, was sufficient to profoundly rescue neuronal maintenance defects in *qv30* null mutant animals. While the rescue is profound, it is not complete, possibly due to the mosaicism of the extrachromosomal array bearing the transgene and the failure to recapitulate normal *sax-7S*(+) expression levels. Nonetheless, larval expression profoundly rescues the null mutants, pointing to the fact that *sax-7S*(+) functions post-developmentally to ensure the maintenance of neuronal organization. This finding rules out the possibility that the neuronal maintenance defects of *sax-7* mutants are a result of an undetected embryonic defect that is amplified by growth and movement of the animal. Instead, our result is consistent with an active requirement for *sax-7* post-embryonically to maintain the organization of an already established nervous system structure.

Thus far, only a handful of molecules have been identified that function to maintain specific aspects of the nervous system. This likely is a reflection of the difficulty associated with determining an adult role for molecules that also play critical roles during development. A post-embryonic neuronal role for *sax-7*, the *C. elegans* homologue of the mammalian L1CAM family, is a conserved property of this gene family. Indeed, loss of L1CAM specifically from the adult mouse brain led to an increase in basal excitatory synaptic transmission and behavioral alterations (Law et al., 2003). In rats, post-developmental nervous system knockdown of Neurofascin severely compromised the already established composition of the axon initial segment and led to an onset of motor deficits (Kriebel et al., 2011; Zonta et al., 2011). Postnatal disruption of CHL1 in excitatory neurons of the mouse forebrain affected the duration of working memory (Kolata et al., 2008). Thus, the continued importance of L1 family members in the adult nervous system is conserved from worm to mammals, suggesting that our findings in *C. elegans* will likely have implications in other organisms.

### SAX-7S is robustly expressed across the nervous system

Transgenic expression of *sax-7S*(+) under different tissue-specific promoters has been used to test for function (this study; (Benard et al., 2012; Diaz-Balzac et al., 2015; Dong et al., 2013; Pocock et al., 2008; Ramirez-Suarez et al., 2019; Salzberg et al., 2013; Sasakura et al., 2005; Zhou et al., 2008; Zhu et al., 2017). Here we have generated *sfgfp* insertion specifically in the *sax-7S* locus, and characterized its endogenous expression pattern. sfGFP::SAX-7S is robustly expressed in virtually all neurons (**Fig. 4**), consistent with the role of SAX-7S in the *C. elegans* nervous system. Indeed, transgenic wild-type copies of *sax-7S*(+) expressed pan-neuronally (P*unc-14::sax-7S, Prab-3::sax-7S*) rescue *sax-7* mutant defects including head ganglia disorganization (**Fig. 2A**), PVQ axon flip-over, AIY and AVK neuronal soma displacement (Pocock et al., 2008), AIY soma position and branching (Diaz-Balzac et al., 2015), AFD neuronal soma position (Sasakura et al., 2005) and PVD length or defasiculation (Ramirez-Suarez et al., 2019), as well as neuronal SAX-7 expression rescues dendrite retrograde extension (Cebul et al., 2020). Interestingly, we observed that sfGFP::SAX-7S expression levels vary among specific neurons in a given animal, and these neuron-specific differences appear to be reproducible across animals. Future studies will address the functional relevance of such SAX-7S expression level signatures.

Transgenic expression of *sax-7S*(+) in the hypodermis (using the epidermal promoter in P*dpy-7::sax-7*(+) transgene), rescues the PVD dendrite defects of *sax-7* mutants (Chen et al., 2019; Dong et al., 2013; Salzberg et al., 2013; Zhu et al., 2017). Despite our careful analyses of animals at all developmental stages, including with unmixing confocal microscopy, we did not observe sfGFP::SAX-7S expression in the body wall epidermis (hyp 7 cells). This suggests that either (1) the endogenous level of SAX-7S in the epidermis is too low to be detected, or (2) the functional form of SAX-7S, in this context, is the C-terminal serine protease cleavage product (fragment B), which cannot be seen with the *qv31* sfGFP::SAX-7S knock-in, as the fluorescent protein is fused N-terminally (**Fig. 5A**). Consistent with this idea, PVD dendrites can be rescued with SAX-7S constructs lacking N-terminal domains Ig3-4 or Ig5-6 (Dong et al., 2013; Salzberg et al., 2013). Tagging the intracellular domain of SAX-7 may allow for visualization of epidermal expression, but such a construct cannot be specific to the short isoform (SAX-7S), if done at the endogenous genomic locus, as both isoforms share the entire intracellular C-terminal region. Previous immuno-histochemistry analyses using an antibody generated against the C-terminal cytoplasmic tail of SAX-7 reported expression of SAX-7 in multiple tissues, including robust signal in neuronal cell bodies, as well as in the nerve ring (major bundle of axons) and the ventral nerve cord (Chen et al., 2001; Wang et al., 2005).

### SAX-7S cleavage products in neuronal maintenance

SAX-7S and SAX-7L proteins could be reliably distinguished on immunoblots thanks to robust controls: the mutant allele *eq1*, where the sequence coding for the intracellular domain of *sax-7* containing the epitope recognized by the antibody is deleted (Chen et al., 2001), and the null *qv30* where the entire *sax-7* locus is deleted. We observed that in wild-type animals, (1) full-length SAX-7S is less abundant than the full-length SAX-7L; (2) the vast majority of SAX-7 protein is cleaved, as an abundant ∼60 kDa cleavage product, seemingly derived from the serine protease-cleavage site; and (3) another less abundant cleavage product of ∼28 kDa may result from cleavage at a site near the transmembrane (**Fig. 1D**). In the *sax-7S*-specific alleles *qv25* and *qv26*, where SAX-7S is absent, the abundance of full-length SAX-7L is similar to wild type, and the ∼60 kDa serine protease cleavage product is less abundant compared to the wild-type, suggesting that the SAX-7S protein may be preferentially cleaved compared to SAX-7L. Also, in the *sax-7L*-specific alleles *eq2* and *nj53*, the ∼60 kDa serine protease-cleavage product appears more abundant than the wild type, perhaps revealing that the SAX-7S cleavage may be favored, resulting in a lower level of full-length SAX-7S versus full-length SAX-7L.

When the serine protease cleavage site is deleted (*sax-7S*-ΔFnIII#3), the resulting recombinant protein is functional in head ganglia maintenance (**Fig. 5B**,**C**), indicating that the cleavage is not essential for function in maintenance of neural architecture, at least with a highly expressed transgene. Consistent with this, motor neuron axon outgrowth defects upon knockdown of *l1camb* in zebrafish can be rescued by expression of a non-cleavable form of L1cam (Linneberg et al., 2019). However, this may be context-specific as Reelin-mediated cleavage of L1CAM in the mouse brain is important for neurodevelopment (Lutz et al., 2017). Furthermore, we find *sax-7S*-ΔFnIII#3 is primarily detected as full-length via western blot (**Fig. 5B**), consistent with recent work which shows that mutating the cleavage site within the third FnIII domain of L1CAM leads to detectable full length protein with no FnIII-domain mediated cleavage products detected (Kleene et al., 2020). This suggests that the third FnIII has conserved importance in SAX-7/L1CAM processing.

Although the two serine protease-cleavage products (SAX-7S-fragment-A and -B) cannot function individually in neuronal maintenance, we find that their simultaneous expression fulfills *sax-7S* neuronal maintenance function. The soluble ectodomain of L1cam similarly cannot solely restore *l1cam* knockdown-mediated defects in motor neuron axon outgrowth in zebrafish L1cam (Linneberg et al., 2019). It is possible that, *in vivo*, serine protease cleavage fragments A and B exist (as suggested by our immunoblot analysis) and may interact together to maintain neuronal architecture. In support of this, furin-mediated cleavage products of Tractin (the L1CAM homologue in leech) can interact *in vitro*, and these fragments together, not individually, can mediate adhesion in an *in vitro* S2 cell aggregation assay (Xu et al., 2003). As we know that SAX-7S can also promote homophilic adhesion in an *in vitro* cell aggregation assay (Sasakura et al., 2005), this points to the intriguing possibility that SAX-7S fragments together *in vivo* may have adhesive neural-maintenance-promoting properties. Future studies will help to address whether these SAX-7S fragments similarly function together or whether their function in neural maintenance is through other interacting factors.

## MATERIALS AND METHODS

### Nematode strains and genetics

Nematode cultures were maintained in an incubator at 20°C (unless otherwise noted) on NGM plates seeded with *Escherichia coli* OP50 bacteria as described (Brenner, 1974). Alleles used in this study are listed in **Table 1**. Strains were constructed using standard genetic procedures and are listed in **Table 2**. Genotypes were confirmed by genotyping PCR or by sequencing when needed. Primers used to build strains are listed in **Table 3**. All the mutant alleles and reporter strains are outcrossed with the Bristol N2 wild-type strain at least 3 times prior to use for analysis or strain building.

**Table 2.**
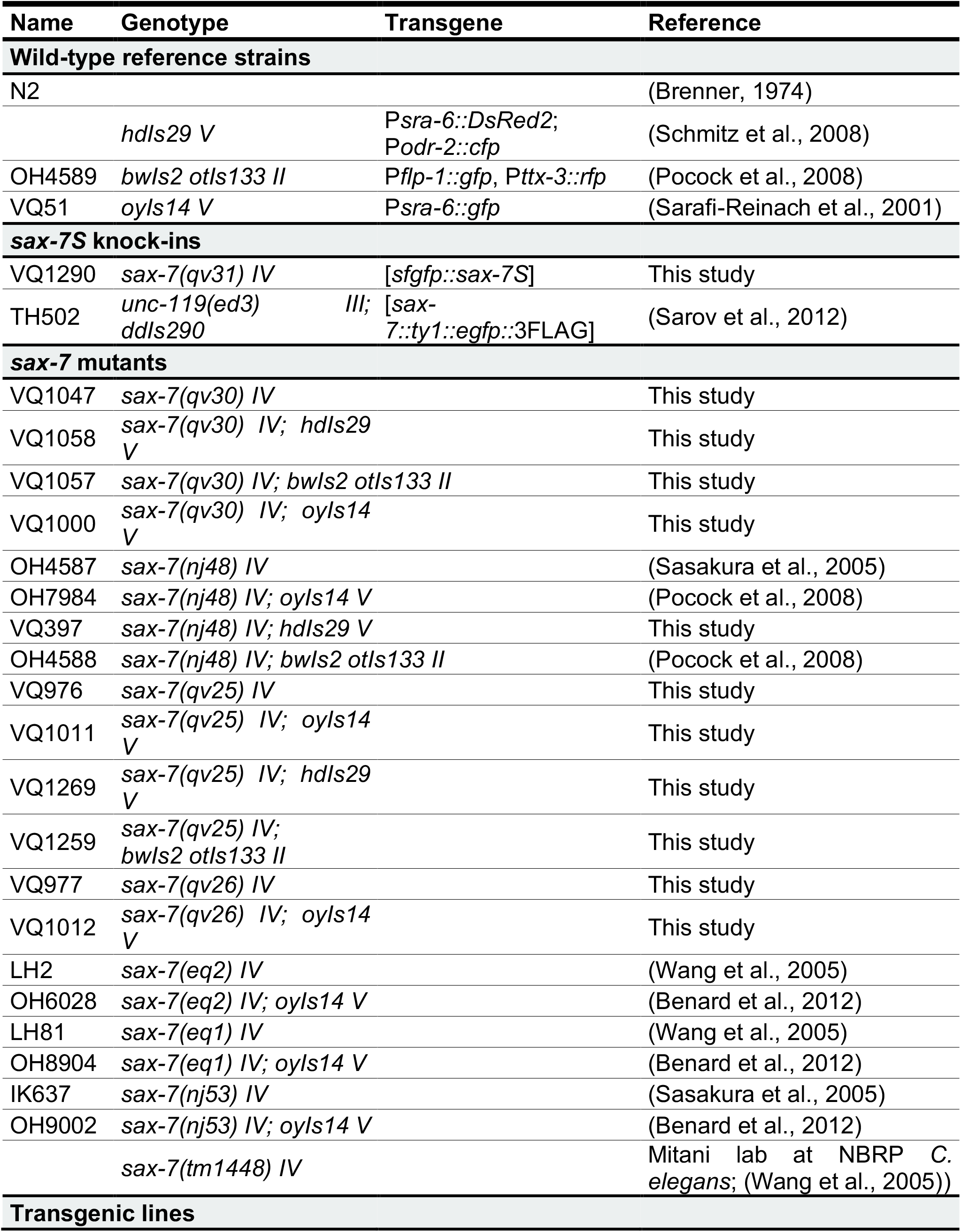
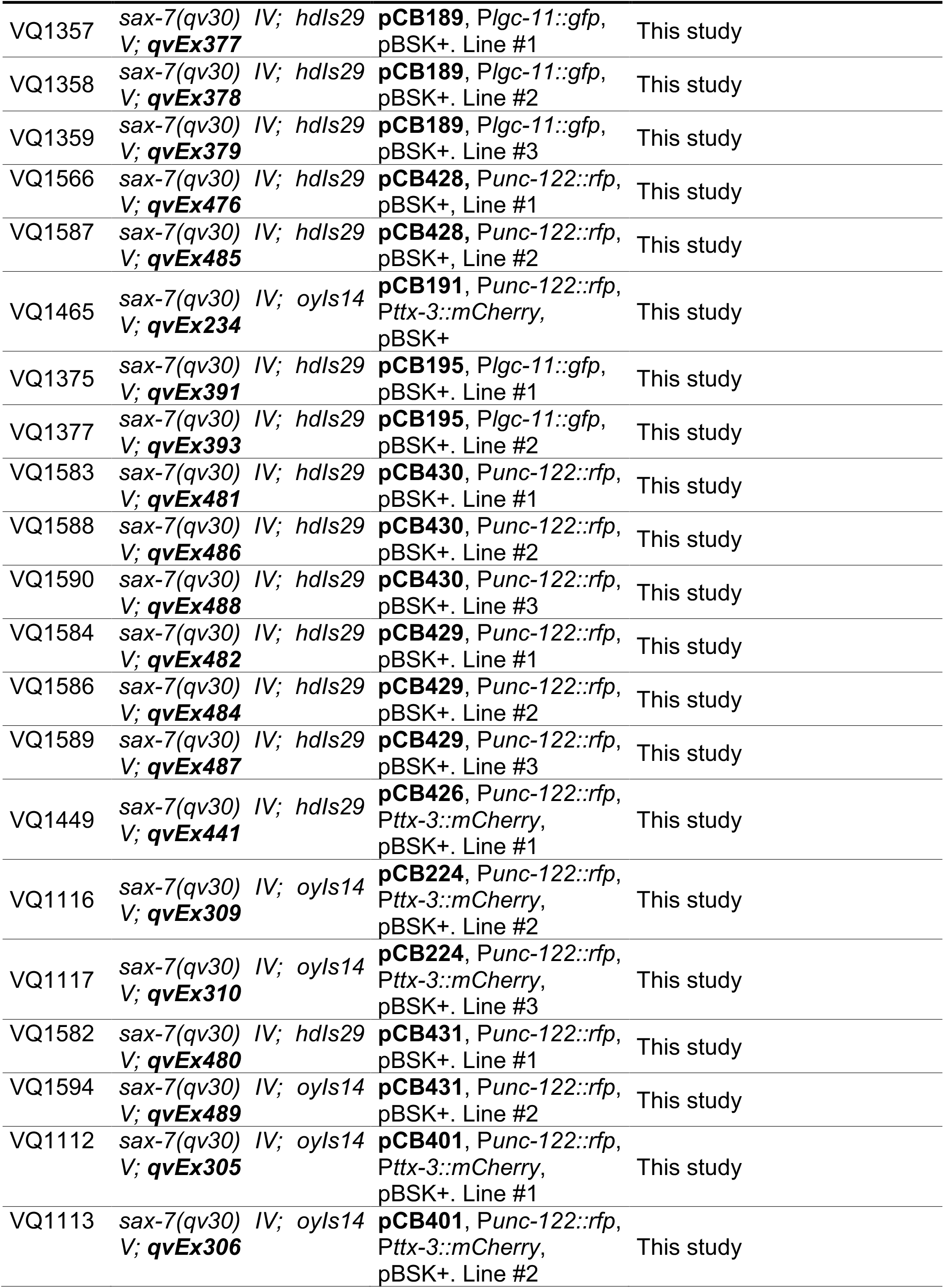
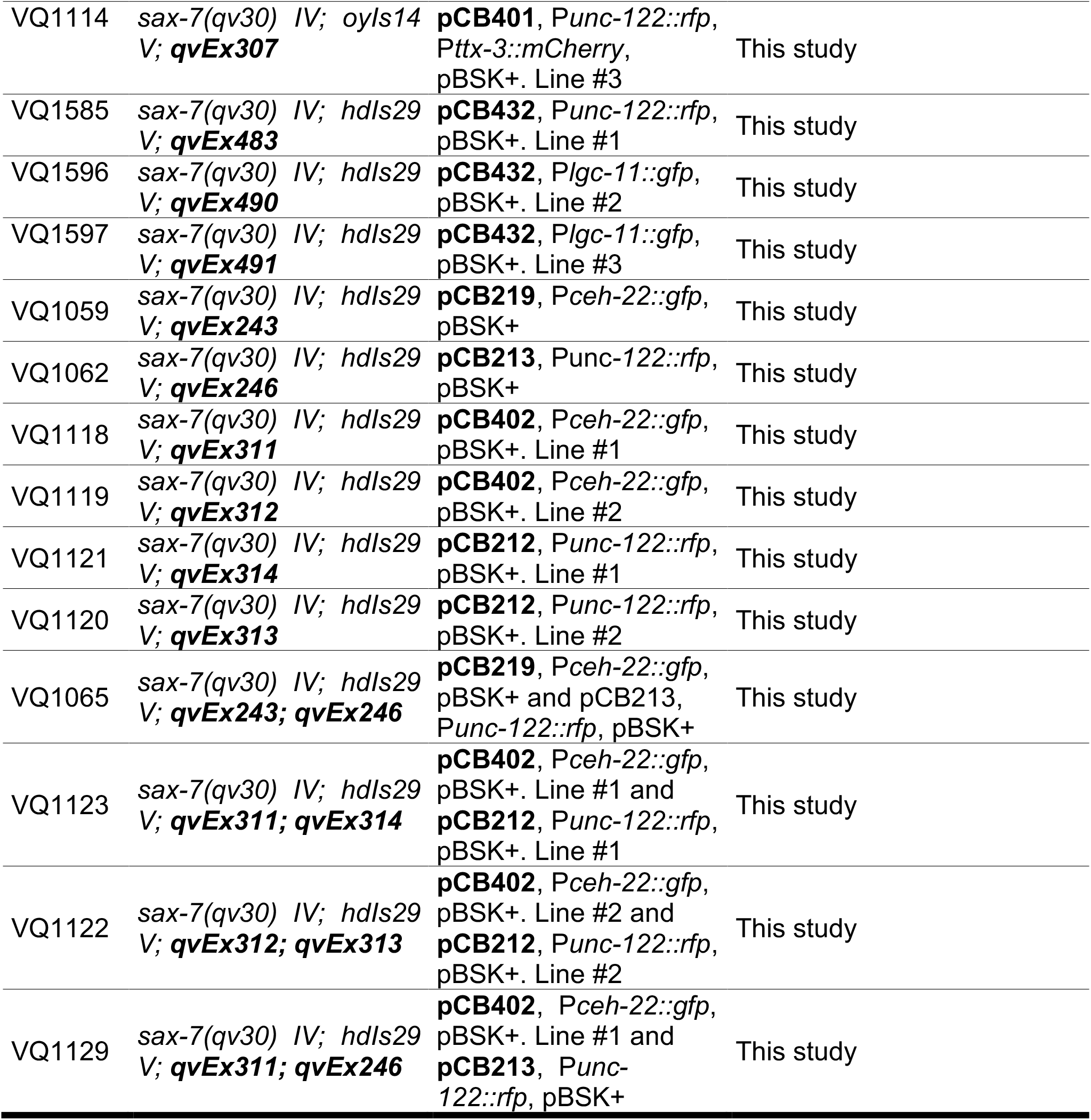
List of strains used.

**Table 3.**
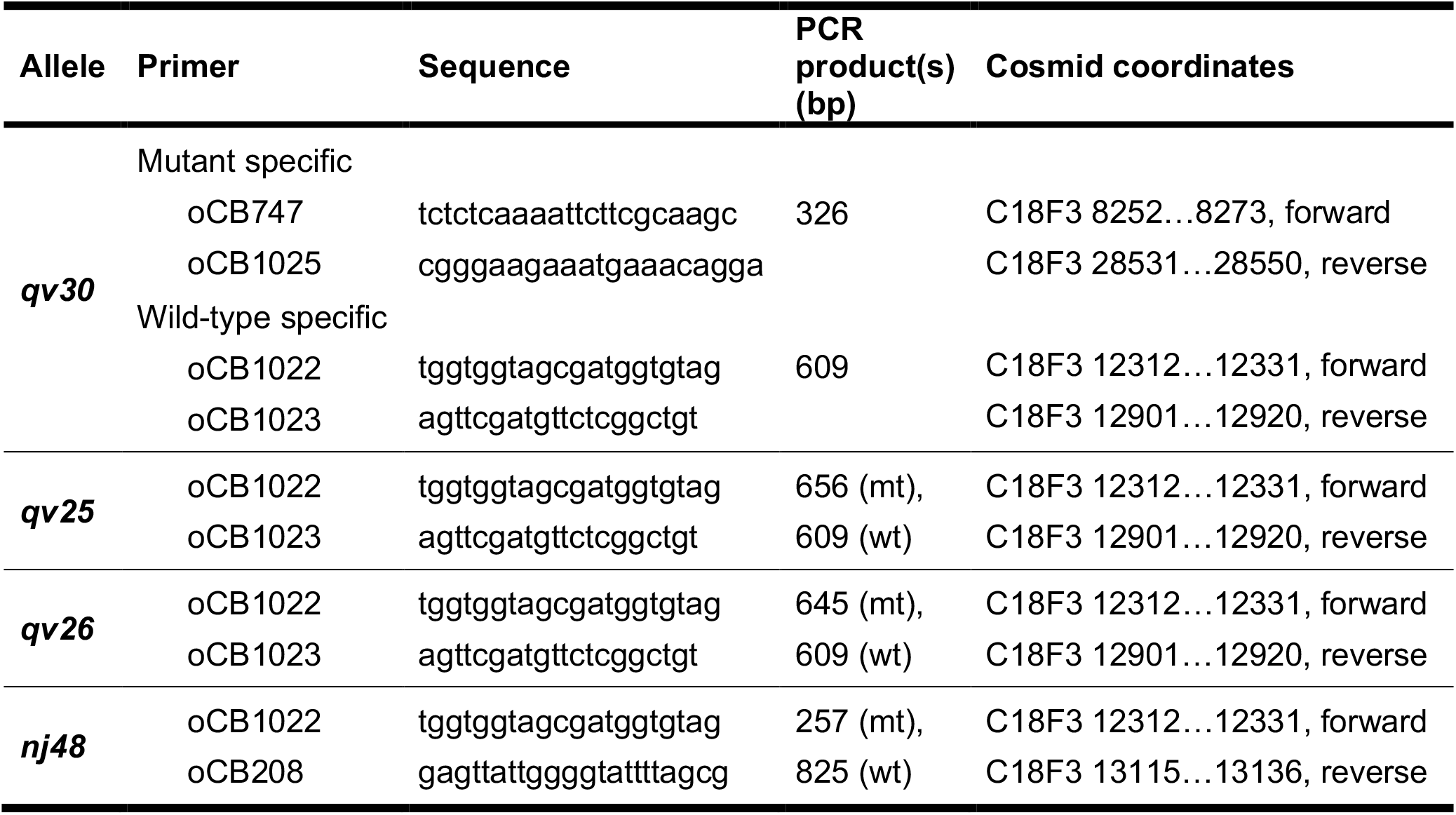
List of primers used to genotype the gene *sax-7* when build strains.

### RT-PCR for *sax-7* alleles

This analysis was performed with wild-type [N2], *sax-7L*-specific mutants [*sax-7(eq2)* and *sax-7(nj53)*], hypomorphic mutants of both isoforms [*sax-7(nj48)* and *sax-7(tm1448)*], and intracellular *sax-7* mutant [*sax-7(eq1)*] strains. Total RNA was extracted from worm samples using Trizol (Invitrogen) according to manufacturer’s instructions. RNA (500 ng) was reverse transcribed using the High Capacity cDNA Reverse Transcription Kit (Applied Biosystems) and random primers. PCR reactions were carried out with 1^st^ strand cDNA template, and 0.25 μM of each primer for *sax-7* cDNA amplification in 10 mM Tris pH 8.3, 1.5 mM MgCl_2_, 50 mM KCl, 0.2 mM deoxynucleotides, and 1 U Phusion DNA polymerase for 30 cycles of 94°C for 10 seconds, 55°C for 20 seconds, and 72°C for 45 secs. Primers used to detect *sax-7* transcript are as following:

oCB985 (CGATTTGCAACTCAACAGGA), oCB986 (TGGTGCTCATGAAGGATCAG), oCB987 (GTGTCCCGAACTGATTCGAT), oCB988 (TTTGTGGAACGTATTGACC), oCB989 (GGAACGTATTGACCTGAAACAG), oCB990 (TTGATCGTCCTGTCCGTGTA), oCB991 (GACCACCGAATACCACAACC).

Primers oCB992 (TCGCTTCAAATCAGTTCAGC) and oCB993 (GCGAGCATTGAACAGTGAAG) were used for the control gene Y45F10D.4 (Hoogewijs et al., 2008) cDNA amplification.

### Generation of *sax-7* null allele by CRISPR-Cas9 (knockout)

#### gRNA plasmids (pCB392 and pCB393)

The gRNAs plasmids were made as previously described (Arribere et al., 2014). To obtain a deletion of the entire locus of *sax-7*, we used two target sequences, one on the 1^st^ exon of the *sax-7* long isoform (gtggccagtgagtaacaag reverse target sequence, pCB392) and the other one on the last exon of *sax-7* corresponding to exon 17 and 14 of long and short isoform, respectively (ccggcatcaagctcttttg reverse target sequence, pCB393).

**pCB392**. Forward and reverse oligonucleotides (oCB1511: AAACcttgttactcactggccacC and oCB1510: TCTTGgtggccagtgagtaacaag, respectively), containing the 5’ target sequence and overhangs compatible with *Bsa*I sites in plasmid pRB1017 (Arribere et al., 2014), were annealed and ligated into pRB1017 cut with *Bsa*I to create the gRNA plasmid pCB392. **pCB393**. Forward and reverse oligonucleotides (oCB1513: AAACcaaaagagcttgatgccggC and oCB1512: TCTTGccggcatcaagctcttttg, respectively), containing the 3’ target sequence and overhangs compatible with *Bsa*I sites in plasmid pRB1017 (Arribere et al., 2014), were annealed and ligated into pRB1017 cut with *Bsa*I to create the gRNA plasmid pCB393.

Plasmids were confirmed by sequencing with M13 reverse primer.

#### The repair donor ssDNA oligonucleotide (repair template)

We designed the repair donor simple-strand DNA oligonucleotide and ordered to Integrated DNA Technologies (IDT) (oCB1514: GATTCTAGATCACGTCGAAAGACCACCATCATGAGGAGCTTCATATTTCTAGCTTGATGCCGGCCGAACGGCCCGAGAAAGGATCAACGTCGACGTTTG, forward). The donor sequence starts with 50 nucleotides corresponding to 5’ homology arm of *sax-7L* at the 5’ target site, followed by 49 nucleotides corresponding to 3’ homology arm of *sax-7* at the 3’ target site.

*sax-7* deletion is located from 8373 bp to 28330 bp on cosmid C18F3, deletion of 19957 bp (**Table 1, Fig. S1A**).

#### Microinjection

DNA mixture was prepared in injection buffer (20 mM potassium phosphate, 3 mM potassium citrate, 2% PEG, pH 7.5). The injection mix contained the Cas9 plasmid (pDD162; (Dickinson et al., 2013) at 50 ng/μL, the gRNA plasmids pCB392 and pCB393 at 50 ng/μL each, the ssDNA donor oCB1514 at 20 ng/μL, the gRNA plasmid pJA58 (*dpy-10* target; (Arribere et al., 2014) at 50 ng/μL and the ssDNA repair template for *dpy-10* (*dpy-10(cn64)*; (Arribere et al., 2014) at 20 ng/μL. Mutations in the *dpy-10* gene were used as CRISPR co-conversion marker.

#### Screening

F1 progeny were screened for Rol and Dpy phenotypes 3-4 days after injection. Rol or Dpy F1 animals were singled and the F2 progeny were screened by PCR for the absence of *sax-7* gene with 2 couples of primers. First couple of primers outside *sax-*7, oCB747 (TCTCTCAAAATTCTTCGCAAGC, forward) and oCB1025 (CGGGAAGAAATGAAACAGGA, reverse), giving a band when *sax-7* is knockout works. Second couple of primers inside *sax-7*, oCB212 (GAAATACACACAAATACGAGTGC, forward) and oCB723 (TAGTTGATTAAAATGTTTCAAGATTG, reverse) giving a band in wild type (no knockout of *sax-7*).

#### Identification

The strain resulting from this genome editing is identified as *sax-7(qv30)* (**Tables 1-3**) and verified by sequencing the deletion junctions (**Fig. S1A**) and also failed to amplify any product by several PCR reactions with primers targeting most of *sax-7* exons.

### Generation of *sax-7(qv25)* and *sax-7(qv26), sax-7S*-specific alleles by CRISPR-Cas9

Two insertion-deletion mutants, namely *sax-7(qv25)* and *sax-7(qv26)* (**Tables 1-3, Fig. S1B-C**), were obtained during our efforts to insert *sfgfp* in the *sax-7S-*specific locus by CRIPSR-Cas9, described below.

### Generation sfGFP::SAX-7S by CRISPR-Cas9 (knock-in)

We chose the protein marker sfGFP as a gene tag because it encodes a GFP variant that folds robustly even when fused to poorly folded proteins and its modified structure resists to the acidic extracellular environment (Pedelacq et al., 2006).

#### gRNA plasmids (pCB394 and pCB395)

The gRNAs plasmids were made as previously described (Arribere et al., 2014). Two target sequences were selected at the end of the exon 1 of *sax-7S-specific* locus (*sax-7S*/C18F3.2a,d), located in the predicted *sax-7S* signal peptide (ggatgtctactgttccttg forward target sequence (pCB394) and tgaaatgaaactaaccaca reverse target sequence (pCB395)).

**pCB394**. Forward and reverse oligonucleotides (oCB1515: TCTTGggatgtctactgttccttg and oCB1516: AAACcaaggaacagtagacatccC, respectively), containing the target sequence and overhangs compatible with BsaI sites in plasmid pRB1017 (Arribere et al., 2014), were annealed and ligated into pRB1017 cut with BsaI to create the gRNA plasmid pCB394. **pCB395**. Forward and reverse oligonucleotides (oCB1518: AAACtgtggttagtttcatttcaC and oCB1517: TCTTGtgaaatgaaactaaccaca, respectively), containing the target sequence and overhangs compatible with BsaI sites in plasmid pRB1017 (Arribere et al., 2014), were annealed and ligated into pRB1017 cut with BsaI to create the gRNA plasmid pCB395.

Plasmids were confirmed by sequencing with M13 reverse primer.

#### The repair donor PCR amplicon (repair template)

We decided to design the repair donor DNA in order that the new gene insertion take place directly at the end of the exon 1 of *sax-7S*, in *sax-7S* signal peptide. The end of the *sax-7S* signal peptide is at beginning of the exon 2 of *sax-7S*. Thus, it was necessary to add this signal sequence part localized downstream the insertion area (TCGGATCGCTACTACACA at the beginning of exon 2) at the end of exon 1, along with the gene *sfgfp* to be inserted, so as to ensure the presence of the entire signal peptide (**Figs. 4B, S1D**).

The repair donor DNA was amplified by PCR using first, primers oCB1525 (GTGTCGGATCGCTACTACACAATGAGCAAAGGAGAAGAAC, forward) and oCB1527 (ATGTGCCCTAAAAAGAAAAATGAAATGAAACTAACTTTGTAGAGCTCATCCATGC, reverse) and a plasmid containing the sequence of sfGFP as template. Primers oCB1525 contains 18 bases in 5’ upstream *sfgfp* corresponding to the missing *sax-7S* signal peptide sequence part and oCB1527 contains 35 bases corresponding to 3’ homology arms of *sax-7S* at the target site. A second PCR was amplified on the previous products with primers oCB1526 (TCATATTCCTGCTAGGATGTCTACTGTTCCTTGTGTCGGATCGCTAC, forward) and oCB1527 (ATGTGCCCTAAAAAGAAAAATGAAATGAAACTAACTTTGTAGAGCTCATCCATGC, reverse). Primer oCB1526 contains 35 bases corresponding to 5’ homology arms of *sax-7S* at the target site. *sfgfp* with signal peptide part were inserted immediately following amino acid 29 of SAX-7S.

#### Microinjection

DNA mixture was prepared in injection buffer (20 mM potassium phosphate, 3 mM potassium citrate, 2% PEG, pH 7.5). The injection mix contained the Cas9 plasmid (pDD162; (Dickinson et al., 2013) at 50 ng/μL, the gRNA plasmids pCB394 and pCB395 at 25 ng/μL each, the *5’arm::sp::sfgfp::3’arm* donor PCR (containing the signal peptide, *sp*) at 100 ng/μL, the gRNA plasmid pJA58 (*dpy-10* target; (Arribere et al., 2014) at 50 ng/μL and the ssDNA repair template for *dpy-10* (*dpy-10(cn64)*; (Arribere et al., 2014) at 20 ng/μL. Mutations in the *dpy-10* gene were used as CRISPR co-conversion marker.

#### Screening

F1 progeny were screened for Rol and Dpy phenotypes 3-4 days after injection. Rol or Dpy F1 animals were singled and the F2 progeny were screened by PCR for the presence of *sax-7S* signal peptide and *sfgfp* in the *sax-7S* locus with primers oCB1022 (TGGTGGTAGCGATGGTGTAG, forward) and oCB818 in *sfgfp* (TTCAGCACGCGTCTTGTAGG, reverse) for the 5’ insertion side and oCB1427 in *sfgfp* (AAAAGCGTGACCACATGGTCC, forward) and oCB1023 (AGTTCGATGTTCTCGGCTGT, reverse) for the 3’ insertion side.

#### Identification

The new strain resulting from this genome editing is identified as *sax-7(qv31[sfgfp::sax-7S])* (**Tables 1, 2**), which is abbreviated as *sfgfp::sax-7S*. The modified locus was verified by sequencing of the entire region (**Fig. S1D**).

### Microinjection to generate transgenic animals

DNA constructs are described in the *Molecular Cloning* section. Briefly, for *sax-7* constructs, the *sax-7* cDNA was subcloned under the control of pan-neuronal promoters *rab-3* (Nonet et al., 1997) and *unc-14* (Ogura et al., 1997) or heat shock promoter *hsp-16*.*2* that express in neurons and other tissues (Fire et al., 1990; Jones et al., 1986; Stringham et al., 1992). Transgenic animals were generated by standard microinjection techniques (Mello and Fire, 1995). Each construct was injected at 1 ng/μL (pCB191), 5 ng/μL (pCB219, pCB213, pCB402 and pCB212), 10 ng/μL (pCB224 and pCB426), or 25 ng/μL (pCB428, pCB189, pCB195, pCB430, pCB429, pCB431, pCB401 and pCB432), along with one or two co-injection markers to select transgenics, including P*ceh-22::gfp* (50 ng/μL) and P*lgc-11::gfp* (50 ng/μL) labelling the pharynx in green, P*ttx-3::mCherry* (50 ng/μL) labelling AIY neurons in red, and P*unc-122::rfp* (50 ng/μL) labelling coelomocytes in red. When needed, pBSK+ was used to increase total DNA concentration of the injection mixes to 200 ng/μL. For details on transgenic strains and their injection mix composition, see **Table 2**.

### Molecular cloning

The gene coding sequences of *sax-7*/C18F3.2b and *sax-7*/C18F3.2a were used for the long and short isoform respectively (available on WormBase). All inserts were verified by sequencing.

#### Construct to express SAX-7S post-developmentally

##### P*hsp16*.*2::sax-7S* (pCB191)

Vector pRP100 (P*unc-14::sax-7S*; (Pocock et al., 2008)) was digested with HindIII and BamHI to release P*unc-14* and ligated with insert of P*hsp-16*.*2* digested out of pPD49.78 (was a gift from Andrew Fire; Addgene plasmid # 1447; RRID: Addgene_1447) with the same restriction enzymes.

#### Constructs to express variants of SAX-7 under pan-neuronal promoters

##### P*rab-3::sax-7S* (pCB428)

*Used for rescue experiments* (Gift from H.E. Bülow (Ramirez-Suarez et al., 2019)).

##### P*unc-14::sax-7S*::Myc (pCB189)

*Used for rescue experiments and western blot against Myc*. Cloned by Gibson assembly. For this plasmid we used the vector pRP100 (P*unc-14::sax-7S*; (Pocock et al., 2008)). The FLAG::*sax-7S*::Myc construct was made by amplifying the 5’ end of the *sax-7S* cDNA from pRP100, carrying a BamHI site, with two nested PCR reactions adding FLAG tag sequence (GATTACAAGGATGACGACGATAAG) right after the signal peptide sequence in the exon 2 and, by amplifying the 3’ end of *sax-7S* cDNA from pRP100, carrying NcoI site, with two nested PCR reactions adding Myc tag sequence (GAGCAGAAACTCATCTCTGAAGAGGATCTG) right before the stop codon, in the exon 14. The vector pRP100 was digested with BamHI and NcoI enzymes to release non-tagged *sax-7S* cDNA in order to clone the synthesized fragment FLAG::*sax-7S*::Myc into it with the same restriction enzymes. As a note, western blot experiments with several anti-FLAG antibodies were done in the attempt of detecting the N-terminus part of SAX-7, but failed.

##### P*unc-14::sax-7L* (pCB195)

*Used for rescue experiments*. Cloned through Gibson assembly. The HA::SAX-7L::V5 construct was made by amplifying the 5’ end of the *sax-7L* cDNA from P*unc-17::sax-7L* construct, carrying BamHI site, with two nested PCR reactions adding HA tag sequence (TACCCATACGACGTCCCAGACTACGCT) after the signal peptide sequence (exon 1) in the exon 2 (between 60-61 *sax-7L* cDNA bases). Also, by amplifying the 3’ end of *sax-7L* cDNA carrying NcoI site, with two nested PCR reactions adding V5 tag sequence (GGTAAGCCTATCCCTAACCCTCTCCTCGGTCTCGATTCTACG) right before the stop codon, in the exon 17. The vector pRP100 was digested with BamHI and NcoI enzymes to release non tagged *sax-7S* cDNA in order to clone the synthesized fragment HA::SAX-7L::V5 into it with the same restriction enzymes.

##### P*unc-14::sax-7S*::Ig3-4 (pCB430)

*Used for rescue experiments* (Gift from H.E. Bülow, (Pocock et al., 2008)).

##### P*unc-14::sax-7S*::Ig5-6 (pCB429)

*Used for rescue experiments* (Gift from H.E. Bülow, (Pocock et al., 2008)).

##### P*unc-14::sax-7S*::FnIII#3 (pCB224)

*Used for rescue experiments* (Pocock et al., 2008).

##### P*unc-14::sax-7S*::FnIII#3::Myc (pCB426)

*Used for rescue experiments and western blot against Myc*. Vector pCB189 (P*unc-14::*FLAG*::sax-7S::*Myc) was digested with PstI and SalI restriction enzymes to release the *sax-7S* cDNA fragment containing the FnIII#3 domain and ligated with insert of *sax-7S* cDNA fragment without the FnIII#3 domain, digested out of pCB224 (P*unc-14::sax-7S::FnIII#3*; (Pocock et al., 2008)) with the same restriction enzymes.

As a note, western blot experiments with several anti-FLAG antibodies were done in the attempt of detecting the N-terminus part of SAX-7, but failed.

##### P*unc-14::sax-7S*ΔFnIII (pCB431)

*Used for rescue experiments* (Gift from H.E. Bülow (Diaz-Balzac et al., 2015)).

##### P*unc-14::sax-7S*ΔAnkyrin (pCB401)

*Used for rescue experiments*. The vector P*ttx-3::sax-7S*ΔAnkyrin (gift from H.E. Bülow) was digested with HindIII and BamHI restriction enzymes to release the fragment P*ttx-3* and ligated with insert of P*unc-14*, digested out of pCB174 (P*unc-14::sax-7L*Δ11, (Pocock et al., 2008)) with the same restriction enzymes.

##### P*unc-14::sax-7S*ΔICD (pCB432)

*Used for rescue experiments* (Gift from H.E. Bülow (Ramirez-Suarez et al., 2019)).

##### P*unc-14::sax-7S* Ig3 to serine protease cleavage site (RWKR) (Fragment A) (pCB219)

*Used for rescue experiments and western blot against Myc*. From pCB189 (P*unc-14::*FLAG*::sax-7S::*Myc), the *sax-7S* cDNA fragment FLAG::Ig3 to serine protease cleavage site (RWKR) (amino acid 745) was amplified with primers oCB798 (CATGATgctagcATGGGGTTACGAGAGACGATGG, forward) and oCB799 (ATCATGccatggCTATCTCTTCCATCTGAACTTTC, reverse) to add on *Nhe*I and NcoI restriction sites, respectively. Vector pCB195 (P*unc-14::*HA*::sax-7L::*V5) was digested with NheI and NcoI and ligated with the insert of *sax-7S* cDNA fragment using the same restriction enzymes. As a note, western blot experiments with several anti-FLAG antibodies were done in the attempt of detecting the N-terminus part of SAX-7, but failed.

##### P*unc-14::sax-7S* serine protease cleavage site (RWKR) to PDZ::Myc (Fragment B) (pCB213)

*Used for rescue experiments and western blot against Myc*. In this case, we needed to be careful adding a signal peptide sequence to assess an accurate expression of the variant. Thus, from pCB189 (P*unc-14::*FLAG*::sax-7S::*Myc), the *sax-7S* cDNA fragment serine protease cleavage site (RWKR) (amino acid 742) to PDZ::Myc was amplified with primers oCB811 (ACTGGCCACATATCATCAGGCAGCATAGATTGGTCAGCGAGATGGAAGAGATCAATTCG, forward) and oCB801 (ATCATGccatggCTACAGATCCTCTTCAGAGATG, reverse) to add the *sax-7L* signal peptide sequence and an NcoI restriction site, respectively. This first nest product was then amplified with primers oCB812 (CATGATgctagcATGAGGAGCTTCATATTCCTCTTGTTACTCACTGGCCACATATCATCAG G, forward) and oCB801 (ATCATGccatggCTACAGATCCTCTTCAG AGATG, reverse) to add NheI restriction site. Then, vector pCB195 (P*unc-14::*HA*::sax-7L::*V5) was digested with NheI and NcoI and ligated with the insert of *sax-7S* cDNA fragment using the same restriction enzymes.

##### P*unc-14::sax-7S* Ig3 to proximal-transmembrane cleavage site (Fragment C) (pCB402)

*Used for rescue experiments*. From pCB189 (P*unc-14::*FLAG*::sax-7S::*Myc), the *sax-7S* cDNA fragment Ig3 to proximal-transmembrane cleavage site (amino acid 1024) was amplified with primers oCB798 (CATGATgctagcATGGGGTTACGAGAGACGATGG, forward) and oCB807 (ATCATGccatggCTAACGAGAACTCGTTCCCGTCG, reverse) to add NheI and NcoI restriction sites, respectively. Then, vector pCB195 (P*unc-14::*HA*::sax-7L::*V5) was digested with NheI and NcoI and ligated with the insert of *sax-7S* cDNA fragment using the same restriction enzymes.

##### P*unc-14::sax-7S* proximal-transmembrane cleavage site to PDZ::Myc (Fragment D) (pCB212)

*Used for rescue experiments*. We needed to be careful adding a signal peptide sequence to assess an accurate expression of the variant. From pCB189 (P*unc-14::*FLAG*::sax-7S::*Myc), the *sax-7S* cDNA fragment from proximal-transmembrane cleavage site (amino acid 1024) to PDZ::Myc was amplified with primers oCB813 (TCACTGGCCACATATCATCAGGCAGCATAGATTGGTCAGCGGAAAGAAATGTCTATCT TTTG, forward) and oCB801 (ATCATGccatggCTACAGATCCTCTT CAGAGATG, reverse) to add the *sax-7L* signal peptide sequence and NcoI restriction site, respectively. This first nested product was then amplified with primers oCB812 (CATGATgctagcATGAGGAGCTTCATATTCCTCTTGTTACTCACTGGCCACATATCATCAG G, forward) and oCB801 (ATCATGccatggCTACAGATCCTCTTCAG AGATG, reverse) to add on NheI restriction site. Then, vector pCB195 (P*unc-14::*HA*::sax-7L::*V5) was digested with NheI and NcoI and ligated with the insert of *sax-7S* cDNA fragment using the same restriction enzymes.

### Protein analysis of endogenous SAX-7 levels in *sax-7* mutant alleles

This analysis was performed with wild-type [*oyIs14*], *sax-7S*-specific mutants [*sax-7(qv25); oyIs14* and *sax-7(qv26); oyIs14*], *sax-7L*-specific mutants [*sax-7(eq2); oyIs14* and *sax-7(nj53); oyIs14*], null mutant [*sax-7(qv30); oyIs14*], hypomorphic mutant of both isoforms [*sax-7(nj48); oyIs14*], and intracellular *sax-7* mutant for antibody specificity control [*sax-7(eq1); oyIs14*] strains. Worms were fed and grown on plates at 20°C for at least three generations before collecting.

For each strain, either (a) 100 L4-stage animals were collected in M9 solution and bacteria was washed off, or (b) large populations of worms were collected. Because the amount of SAX-7 protein was too low to detect all the protein forms on the analysis above, large pellets of thousands of mixed-stage worm populations were collected by washing plates, mostly devoid of bacteria, with M9 solution.

NETI (NaCl, EDTA, Tris, IGEPAL) buffer and protease inhibitors (Roche #11836153001) were added to worm pellets with 2X Laemmli sample buffer (Bio-Rad #161-0737) and 5% β-mercaptoethanol (v/v), and immediately frozen in liquid nitrogen. Samples were boiled for 5 min at 95°C and centrifuged for 10 min at 10000 rpm prior to loading with capillary tips. Proteins were separated by SDS-PGE on a 4-15% Mini-PROTEAN^®^ TGX Stain-Free™ gel (Bio-Rad #456-8084) and transferred with the Trans-Blot^®^ Turbo™ RTA Transfer Kit (Bio-Rad #170-4275) to a LF (low fluorescence) PVDF membrane using the Trans-Blot^®^ Turbo™ Transfer System (Bio-Rad). Membranes were blocked in 5% BSA (VWR #0175), 5% non-fat milk and incubated in 1:8000 rabbit anti-SAX-7, an affinity purified antibody generated against the SAX-7 cytoplasmic tail [gift of (Chen et al., 2001)] and 1:5000 goat anti-rabbit HRP secondary antibody (Bio-Rad #170-5046). For the loading control, membranes were incubated in 1:1000 rabbit anti-HSP90 antibody (CST #4874) and 1:5000 goat anti-rabbit HRP secondary antibody (Bio-Rad #170-5046). Signal was revealed using Clarity Max™ Western ECL Substrate (Bio-Rad #170-5062), and imaged using the ChemiDoc™ System (Bio-Rad). This analysis was performed three times for each set of experiments.

### Protein analysis of transgenic SAX-7S protein fragments

Myc tag was used (see *sax-7S* transgenes section). This analysis was performed with wild-type [*hdIs29*] and *qv30* null mutant transgenic animals carrying different *sax-7S* protein fragments under the pan-neuronal promoter P*unc-14*: *sax-7S* “Fragment A” [Ig3 up to serine protease cleavage site] (VQ1059), “Fragment B” [serine protease cleavage site up to C-terminal::Myc] (VQ1062), *sax-7S-*A and -B fragments together [Ig3 up to serine protease cleavage site and serine protease cleavage site up to C-terminal::Myc] (VQ1065), *sax-7S* full length [*sax-7S*::Myc] (VQ1357), and *sax-7S* without serine protease cleavage site in the 3^rd^ FnIII [*sax-7S*ΔFnIII#3::Myc] (VQ1449).

For each strain, before collecting, worms were allowed to grow for ∼2 generations by feeding with ∼30 transgenic worms (or non-transgenic for wild type). Because these assays require large pellets of thousands of worms, rather than picking transgenic animals, worms were collected by washing populations on plates. We estimate that around ∼50% of animals carry the various extrachromosomal transgenes (described above). Indeed, unstable non-integrated extrachromosomal arrays are lost during cell divisions and over generations, so that by the time that the worms were collected from plates, not all, but a proportion of the animals on the plates are transgenics (this was verified by visual inspection). Mixed-stage worm populations from plates devoid of bacteria were collected in M9 solution. Then, NETI (NaCl, EDTA, Tris, IGEPAL) buffer and protease inhibitors (Roche #11836153001) were add to worm pellets with 2X Laemmli sample buffer (Bio-Rad #161-0737), 5% β-mercaptoethanol (v/v), and immediately frozen in liquid nitrogen. Each sample provided enough material to load 2 gel wells allowing the visualization of SAX-7S recombinants tagged by Myc. Samples were boiled for 5 min at 95°C and centrifuged for 10 min at 10000 rpm prior to loading with capillary tips, separated by SDS-PGE on a 4-15% Mini-PROTEAN^®^ TGX Stain-Free™ gel (Bio-Rad #456-8084), and transferred with the Trans-Blot^®^ Turbo™ RTA Transfer Kit (Bio-Rad #170-4275) to a LF (low fluorescence) PVDF membrane using the Trans-Blot^®^ Turbo™ Transfer System (Bio-Rad). Membranes were blocked in 5% BSA (VWR #0175), 5% non-fat milk. Blots were incubated in 1:500 mouse anti-Myc (CST #2276) and 1:3000 goat anti-mouse HRP secondary antibody (Jackson ImmunoResearch #115-035-003). For the loading control, membranes were incubated in 1:1000 rabbit anti-HSP90 antibody (CST #4874) and 1:5000 goat anti-rabbit HRP secondary antibody (Bio-Rad #170-5046). Signal was revealed using Clarity Max™ Western ECL Substrate (Bio-Rad #170-5062), and imaged using the ChemiDoc™ System (Bio-Rad). This analysis was performed three times.

### Microscopy and imaging

Worms were grown in incubator at 20°C for at least 3 generations prior to analysis. Worm stages are indicated in the figures. 24 h post-L4 stage is considered “1^st^ day of adulthood”, 24 h after that is considered “day 2 of adulthood”, and so on.

### Neuroanatomical observations

Neuroanatomy was examined in wild-type and mutant animals using specific reporters. Worms were anesthetized with 75 mM of sodium azide (NaN_3_) and mounted on 5% agarose pads on glass slides. Animals were observed with Nomarski or fluorescence microscopy (Carl Zeiss Axio Scope.A1 or Axio Imager.M2), and images were acquired using the AxioCam camera (Zeiss) and processed using AxioVision (Zeiss), with 60x oil immersion objective (expected for PVQ/PVP axons: 100x oil immersion objective).

#### Analysis of ASH/ASI cell body positioning with respect to the nerve ring

Cell body pairs of ASH/ASI chemosensory neurons and the nerve ring (neuropil of the worm), positioned in the head ganglia of the worm, were visualized using *hdIs29* (Schmitz et al., 2008), an integrated P*sra-6::DsRed2*; P*odr-2::cfp* reporter as well as *oyIs14 (Sarafi-Reinach et al*., *2001)*, an integrated P*sra-6::gfp* reporter. Animals were analyzed in a lateral orientation. Normally, both the two ASH and the two ASI soma are located posterior to the nerve ring. Animals were counted as mutant when at least one ASH or ASI soma was touching, on top of, or anterior to the nerve ring. Animals were counted as wild type when all ASH/ASI soma were positioned posterior to the nerve ring.

#### Analysis of AVK/AIY soma

Cell body pairs of AVK/AIY interneurons, were visualized using a stock containing two integrated reporters, *bwIs2* (P*flp-1::gfp*) to label AVK in green, and *otIs133* (P*ttx-3::rfp*) to label AIY in red (Pocock et al., 2008). Animals were analyzed when in a ventral orientation. Cell bodies of AVK/AIY localized to the head ganglia of the worm in the retrovesicular ganglion. Normally, both neuron pairs AVKL/AIYL (left) and AVKR/AIYR (right) adhere to each other (Pocock et al., 2008; White et al., 1986b). Animals were counted as mutant when one or two AVK/AIY pairs were detached. Animals were counted as wild type when both of the AVK/AIY soma pairs were in contact.

#### Analysis of PHA/PHB soma

Cell body pairs of PHA/PHB chemosensory neurons, were visualized using DiI (1,1’-Dioctadecyl-3,3,3’,3’-Tetramethylindocarbocyanine Perchlorate) staining procedure (Hedgecock et al., 1985). This is a lipophilic fluorescent stain for labeling cell membranes and hydrophobic structures, providing an alternative for labeling cells and tissues. In our case, it allows us to stain and visualize by a pink fluorescence the ciliated amphid (ADL, ASH, ASI, ASJ, ASK, AWB) and phasmid (PHA, PHB) neurons (Collet et al., 1998), that are exposed to the outside environment. Animals were analyzed in a ventral orientation. Cell bodies of PHA/PHB localized to the tail ganglia of the worm in the lumbar ganglion. Normally, both neuron pairs PHAL/PHBL (left) and PHAR/PHBR (right) adhere to each other (White et al., 1986a). Animals were counted as mutant when any of the PHA/PHB pairs were detached from one another. Animals were counted as wild type when both of the PHA/PHB soma pairs were in contact.

#### Analysis of PVQ/PVP axons

PVQ and PVP axons were visualized in animals using *hdIs29* (Schmitz et al., 2008), an integrated P*sra-6::DsRed2*; P*odr-2::cfp* reporter, labelling PVQ and PVP in red and green, respectively. Animals were analyzed in a ventral orientation. The axons of the PVQL/PVPL and the PVQR/PVPR neurons are normally located within the left and right fascicle of the ventral nerve cord, respectively. Animals were counted as having an axon flip-over defect when one of the PVQ/PVP axons was flipped to the opposite fascicle at any point along the ventral nerve cord, as previously described (Benard et al., 2006).

### Other phenotypic observations

#### Analysis of embryonic lethality

From plates with hermaphrodites laying eggs, a lot of embryos were picked with OP50, and spread into a new plate that was kept at 20°C. After β16 hr, the number of larvae and dead embryos were counted. This experiment was repeated 3 times. *Analysis of brood size*. An L4 worm was singled on a new plate independently. The number of embryos laid were counted each day of adulthood until 4-days-old adults and the total amount of laid embryos during 4 days was calculated. This was done at least 7 times.

#### Analysis of egg-laying

Ten L4-stage worms were put on one plate and their ability to lay embryos normally was examined each day from day 1 to 5 of adulthood. Worms deficient in embryo laying retain them inside their bodies and display an Egl phenotype (Desai and Horvitz, 1989; Trent et al., 1983). When counted defective they were removed from the plate. This was done 10 times.

### Expression pattern analysis of sfGFP::SAX-7S

Fluorescence images of *sax-7::ty1::egfp::*3FLAG strain (**Fig. 4A**; (Sarov et al., 2012)) were captured by fluorescence microscopy (Carl Zeiss Axio Imager.M2), and images were acquired using the AxioCam camera (Zeiss) and processed using AxioVision (Zeiss), with 60x oil immersion objective.

Fluorescent images of *qv31*, the *sfgfp::sax-7S* strain (**Fig. 4B**), were captured using a Nikon A1R confocal microscope and processed using ImageJ. For each stage, at least 8 worms were examined in detail. Nematodes were immobilized in 75 mM of NaN_3_ and mounted on 5% agarose pads on glass slides. All fluorescence images for *sfgfp::sax-7S* strain were obtained with the same settings using a Nikon Ti-e spinning disk confocal with 60x oil immersion objective. Images were three-dimensionally unmixed with NIS-Elements image acquisition and analysis software. Green fluorescent background is commonly seen in worms (gut granules), which disturbs the analysis of green fluorescent fusion proteins. In this study, we took advantage of a microscopy technique which “unmixes” overlapping spectral emissions after acquisition. Thanks to highly sensitive GaAsP-detectors, signals can be distinguished by the process called “spectral unmixing” (Ackermann, 2017).

For this, we acquired images for wild type N2 animals and determined a ROI in the pharynx in the head of the worm, giving a spectral profile defined as “background” green auto-fluorescence the worm. Then, with the *sfgfp::sax-7S* CRISPR-Cas9 strain, which expresses “real” green fluorescence, we acquired images and determined a ROI to the soma part of one neuron in the head of the worm, giving a spectral profile defined as “real” green fluorescence in the case of the sfGFP fluorophore. Finally, the “background” profile was subtracted from the “real” green fluorescence profile, keeping the real green fluorescence emission coming from sfGFP for the entire animal. ND2 files generated with NIS-Elements were imported into Fiji for analysis. Maximum intensity projections were generated by selecting stacks that had both ventral and dorsal signals.

### Heat-shock inducible expression of *sax-7S*(+)

This analysis was performed with wild type [*oyIs14*], null mutant [*sax-7(qv30); oyIs14*] and null mutant transgenic animals carrying *sax-7S* cDNA under heat shock promoter *hsp16*.*2* that express in neurons and other tissues (Fire et al., 1990; Jones et al., 1986; Stringham et al., 1992). Worms were maintained in the incubator at 15°C for at least two generations prior to analysis. To generate freshly hatched pools of L1s, plates were fed with a lot of adult hermaphrodites (which carry eggs) and left at 15°C around 15h (overnight) in order to have many laid embryos close to hatch. Then, embryos were picked on a new plate and kept for β6h at 15°C, after which any remaining unhatched embryos were removed from the plates leaving only freshly hatched L1s (on average 3.5 h old) on the plate. Animals were either heat shocked immediately as freshly L1s, or as L3s (∼42 h post-hatch). Heat shock treatment consisted of 3 cycles of 30 minutes at 37°C with a 60 minutes recovery period at 20°C between each cycle, after which plates were put back at 15°C until analysis as adults (**Fig. 3A**). All experiments were repeated at least twice. Neuroanatomical analysis of ASH/ASI cell body positioning with respect to the nerve ring (see Neuroanatomical observations) were performed on animals as 1-, 2-, 3-, 4-, and 5-days-old adults.

### Quantification and statistical analysis

z-tests and student’s t test were performed in MS Office Excel. Error bars in bar graphs represent standard error of proportion (S.E.P.).

## Data availability

Mutant and genome engineered strains will be available at the *Caenorhabditis Genetics Center*, and all strains and plasmids are available upon request. The authors affirm that all data necessary for confirming the conclusions of the article are present within the article, figures, and tables.

## ACKNOWLEDGMENTS

We thank Maria Doitsidou and Lise Rivollet for comments on the manuscript; Denis Flipo for assistance with confocal microscopy and unmixing; Andrea Thackeray and Lise Rivollet for help throughout the project; the following researchers for sharing reagents: Max Heiman (for a plasmid containing the *sfgfp* gene); Lihsia Chen (for anti-C-terminal SAX-7 antibodies); Roger Pocock and Hannes Bülow for plasmids, as well as the CGC, which is funded by the NIH Office of Research Infrastructure Programs (P40 OD010440), and WormBase. This work was supported by funds from the NIH, CIHR, NSERC, and FRQS to C.Y.B.; and scholarships to V.E.D. by UQÁM and the CERMO-FC Research Center.

## SUPPLEMENTARY FIGURE LEGENDS

**Fig S1.**
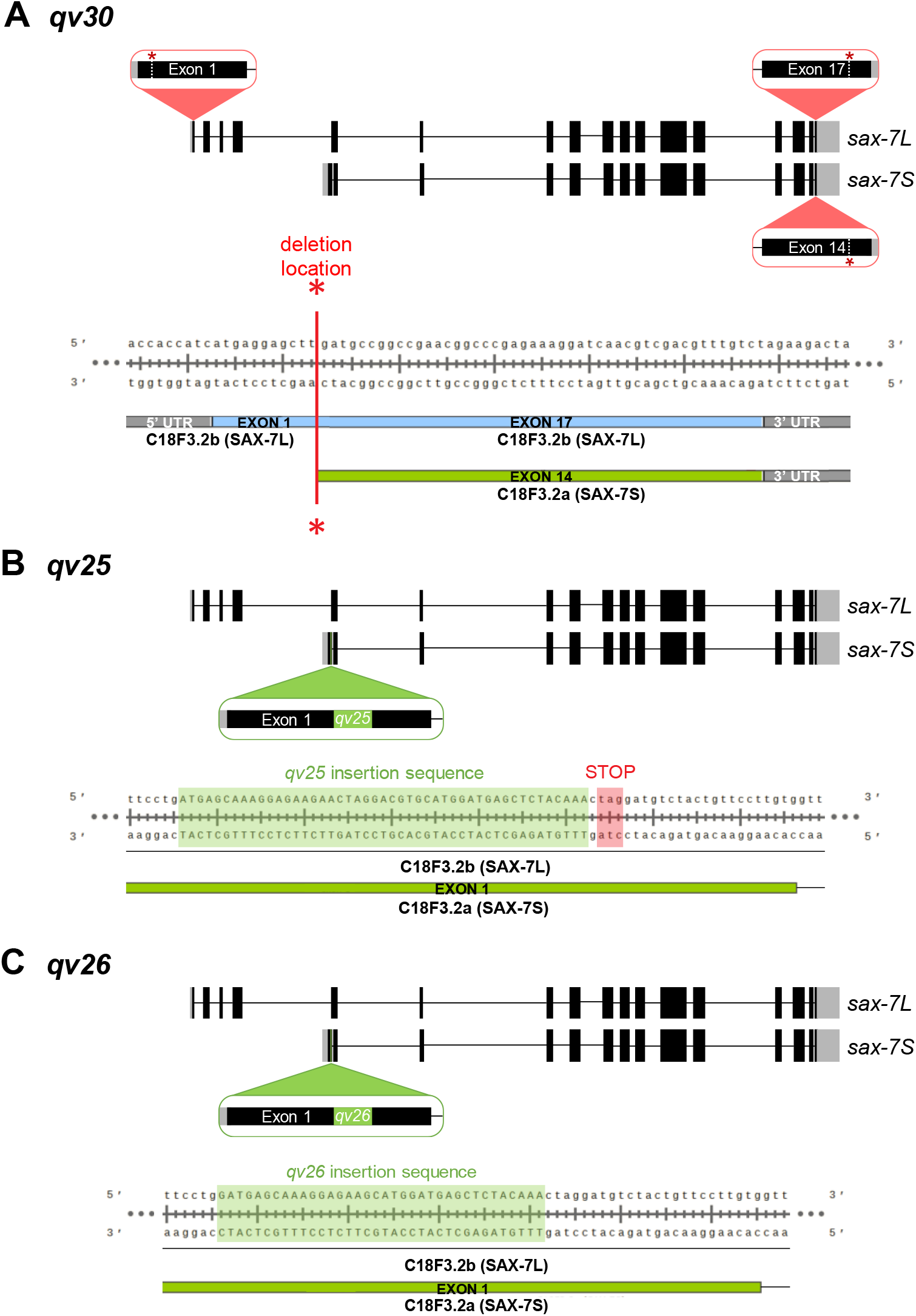
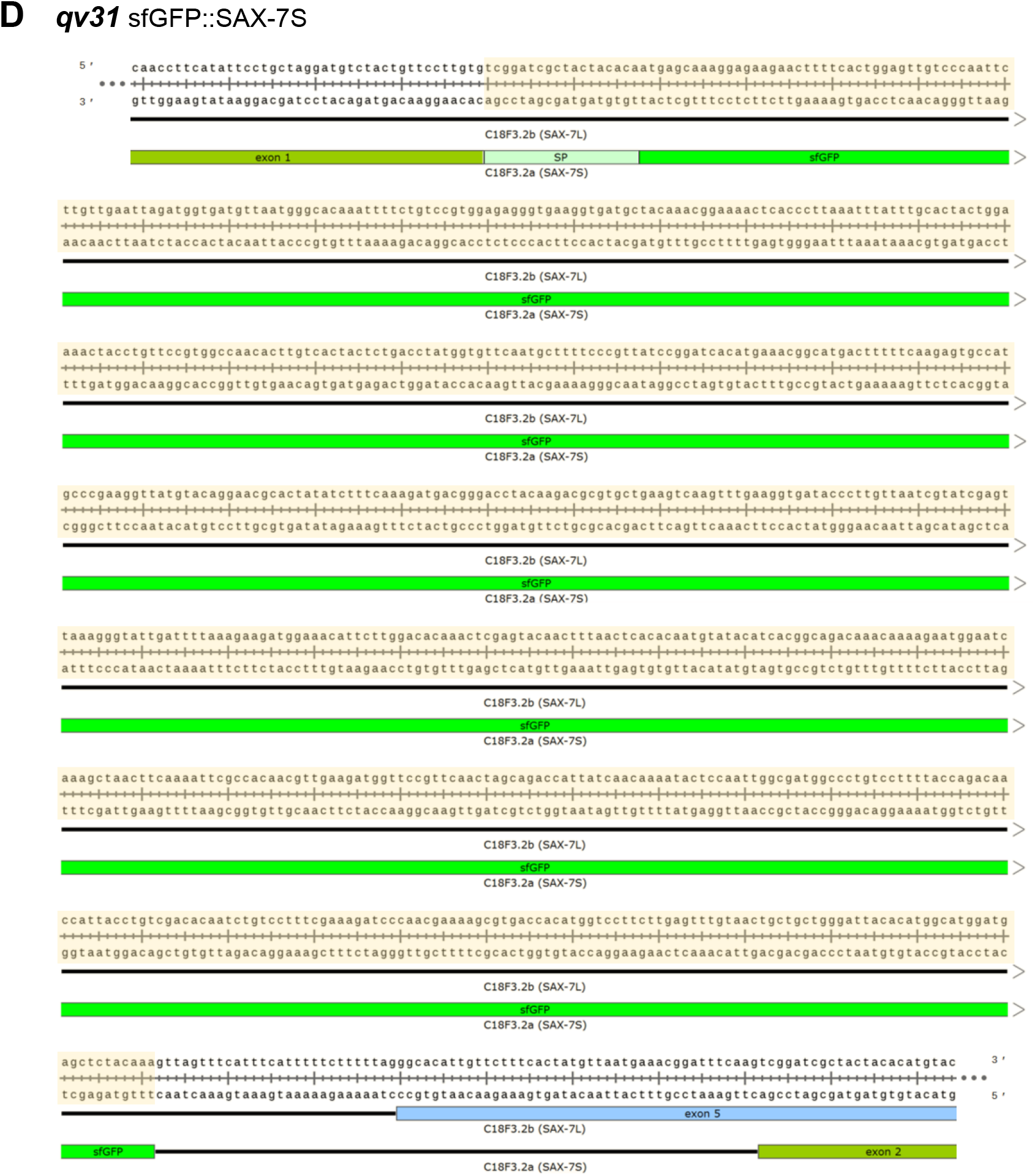
Sequence information for the four CRISPR-Cas9 genome edited alleles generated in this study. **(A)** Diagram of the *qv30* allele. It is a 19973 bp deletion, from base 8364 in exon 1 of *sax-7L* on cosmid C18F3 (indicated by white dotted line with a red asterisk in exon 1) to base 28337 (indicated by white dotted line with red asterisk in exon 17 of *sax-7L*, and exon 14 of *sax-7S*). Below, the exact sequence context of the *qv30* deletion is provided (with the red line and asterisks), on what remains of *sax-7L* exons in blue, and of *sax-7S* exons in green. Exons of *sax-7L* and *sax-7S* are represented by blue and green boxes, respectively, and UTRs by grey boxes. **(B)** Diagram of the *qv25* allele. It is an insertion of 47 bp (indicated in green), right after bp 12785 exon 1 of *sax-7S* on C18F3. It creates an ORF frameshift and introduces a stop codon in the *sax-7S* signal peptide (the signal peptide is encoded by sequence on exon 1 and beginning part of exon 2). *sax-7S* exons are represented by green boxes and introns by black lines. **(C)** Diagram of the *qv26* allele. It is an insertion of 36 bp (indicated in green), right after bp 12785 exon 1 of *sax-7S* on C18F3. It disrupts the *sax-7S* signal peptide (the signal peptide is encoded by sequence on exon 1 and beginning part of exon 2). *sax-7S* exons are represented by green boxes and introns by black lines. **(D)** Diagram of the *qv31* allele. In order to tag the endogenous short isoform (*sax-7S*) specifically, the gene coding for superfolder GFP (*sfgfp*, 732 bp), preceded by the downstream part of the coding sequence for the *sax-7S* signal peptide (from beginning exon 2 of *sax-7S*), was inserted by CRISPR-Cas9-mediated homology repair. This insertion (highlighted in yellow) starts right at the end of exon 1 of *sax-7S*, after bp 12809 of C18F3. “SP” means *sax-7S* signal peptide sequence part added. Exons of *sax-7L* and *sax-7S* are represented by blue and green boxes, respectively, and introns by black lines.

**Fig S2.**
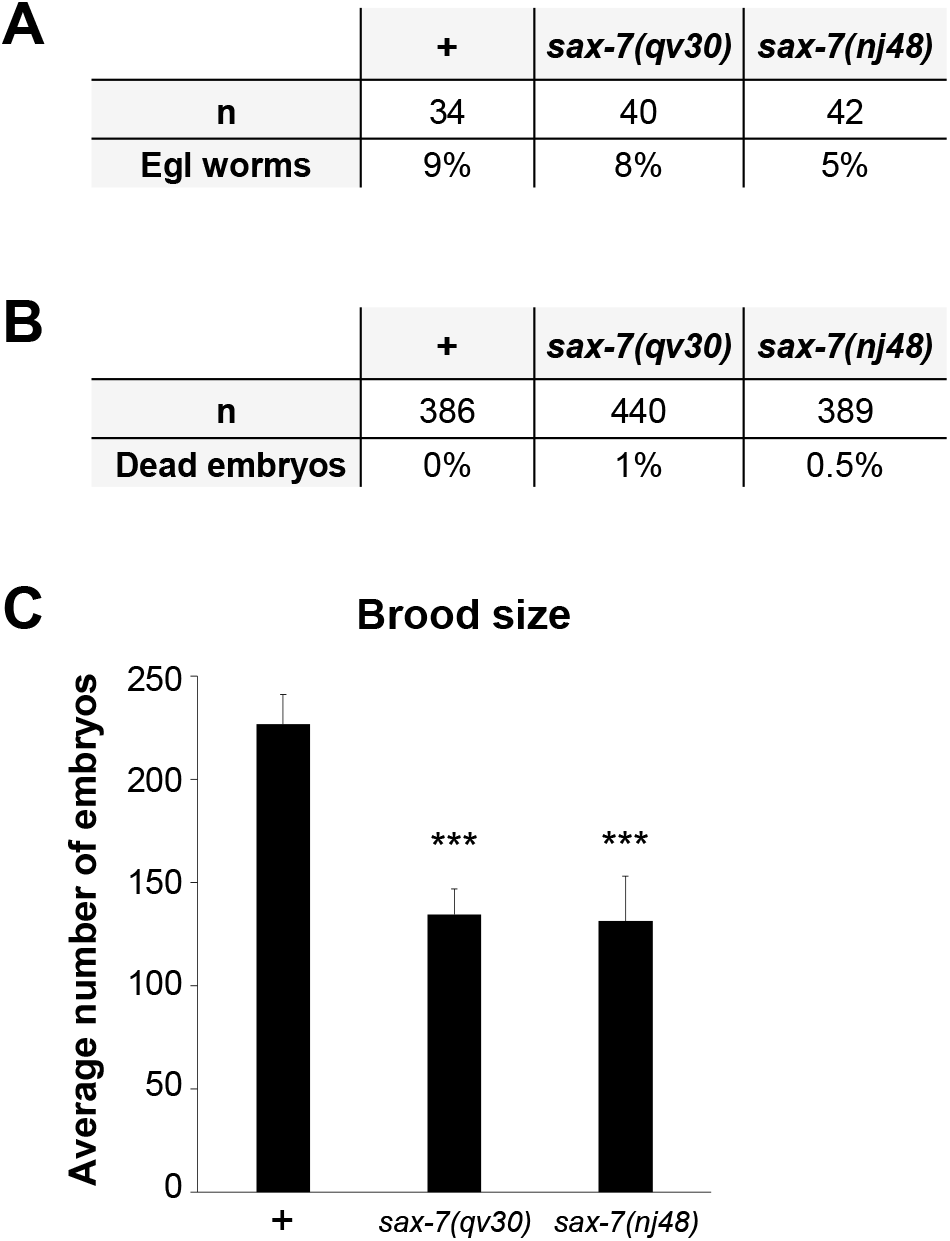
Phenotypic characterization of *sax-7(qv30)*. **(A)** *sax-7* mutants have normal egg laying behavior. Ability to normally lay embryos was examined each day from day 1 to 5 of adulthood. **(B)** *sax-7* mutants have normal embryonic viability. **(C)** *sax-7* mutants have a smaller brood size. Quantification of the total number of embryos laid from L4 until 4-day-old in wild type, null mutant *qv30*, and hypomorphic mutant *nj48*.

**Fig S3.**
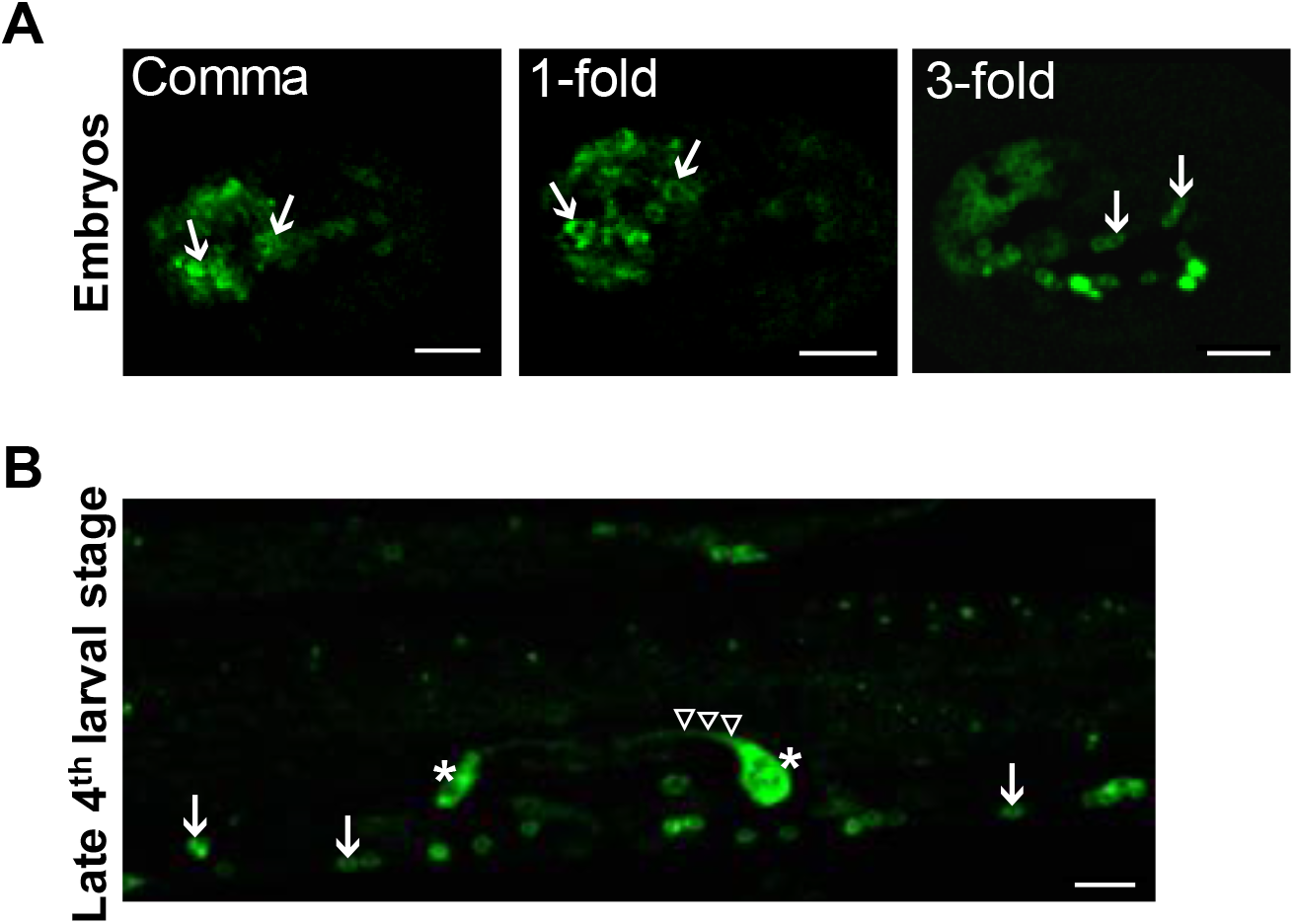
Other sites of SAX-7S expression. **(A)** Unmixed confocal images showing sfGFP::SAX-7S expression (green fluorescence) in neurons at different embryonic stages. No fluorescence was observed in 28-cell stage embryos (data not shown). White arrows indicate sfGFP::SAX-7S expression in embryonic neurons, localized to the plasma membrane. **(B)** Unmixed confocal images showing sfGFP::SAX-7S expression (green fluorescence) in the developing reproductive system in late 4^th^ larval stage uterus. sfGFP::SAX-7S expression is seen in the utse syncytium (empty white arrowhead), in two uterine ventral cells (likely uv1, white asterisks) and in neurons of the ventral nerve cord (white arrows) of the worm. n ≥ 20 animals examined by confocal microscopy for each stage. z-stack maximum intensity projections. Scale bar, 10 μm.

